# MAPK and mTORC1 signaling converge on Cyclin D to enable cell-cycle re-entry in melanoma persister cells

**DOI:** 10.1101/2025.05.14.653869

**Authors:** Varuna Nangia, Humza Ashraf, Nasreen Marikar, Victor J. Passanisi, R. Christopher, Sabrina L. Spencer

## Abstract

Resistance to targeted therapy in BRAF-mutant melanomas can arise from drug-tolerant persister cells, which can non-genetically escape drug-induced quiescence to resume proliferation. To investigate how melanoma cells escape drug to re-enter the cell cycle within 3-4 days of BRAF and MEK inhibition, we computationally reconstructed single-cell lineages from time-lapse imaging data, linking key signaling pathways to cell-cycle fate outcomes. We found that ERK reactivation, while necessary, is insufficient for cell-cycle re-entry under MAPK inhibition. Instead, mTORC1 emerged as an additional critical mediator, enhancing translation and cell growth in cells destined for drug escape. We further found that ERK and mTORC1 signaling converge to produce Cyclin D1 protein levels, a key bottleneck for cell-cycle re-entry. Using CRISPR to tag endogenous Cyclin D1, we found that future escapees markedly upregulate Cyclin D1 at least 15 hours prior to drug escape, compared to non-escapees. Importantly, this early upregulation enables accurate prediction of future escapees from non-escapees, underscoring how differences in Cyclin D1 accumulation precede and govern the timing and likelihood of cell-cycle re-entry in persister cells. Our findings suggest that variability in ERK and mTORC1 activity underlies the heterogeneous Cyclin D1 levels observed, influencing the proliferative potential of persister cells and ultimately shaping the diverse cell-cycle behaviors observed under drug treatment. Cyclin D1 protein therefore emerges as both a critical biomarker and a therapeutic target for preventing cell-cycle re-entry in BRAF/MEK-treated melanoma.

**One Sentence Summary:** Cyclin D1 accumulation, driven by the integration of heterogeneous MAPK and mTORC1 signaling, is a critical bottleneck for cell-cycle re-entry in drug-treated melanoma cells.

## INTRODUCTION

Cell proliferation is regulated by the mitogen-activated protein kinase (MAPK) pathway, which relays extracellular mitogen signals from receptor-tyrosine kinases (RTK) through the canonical RAS-RAF-MEK-ERK signaling cascade. Once active, ERK stimulates a family of transcription factors that upregulate D-type cyclins, which in turn activate Cyclin-Dependent Kinases (CDK) 4 and 6 (*1*). CDK4/6 then phosphorylates and inactivates the retinoblastoma tumor suppressor, Rb, triggering a positive feedback loop that promotes cell-cycle entry (*2*) (Fig. 1A, *left*).

**Figure. 1.**
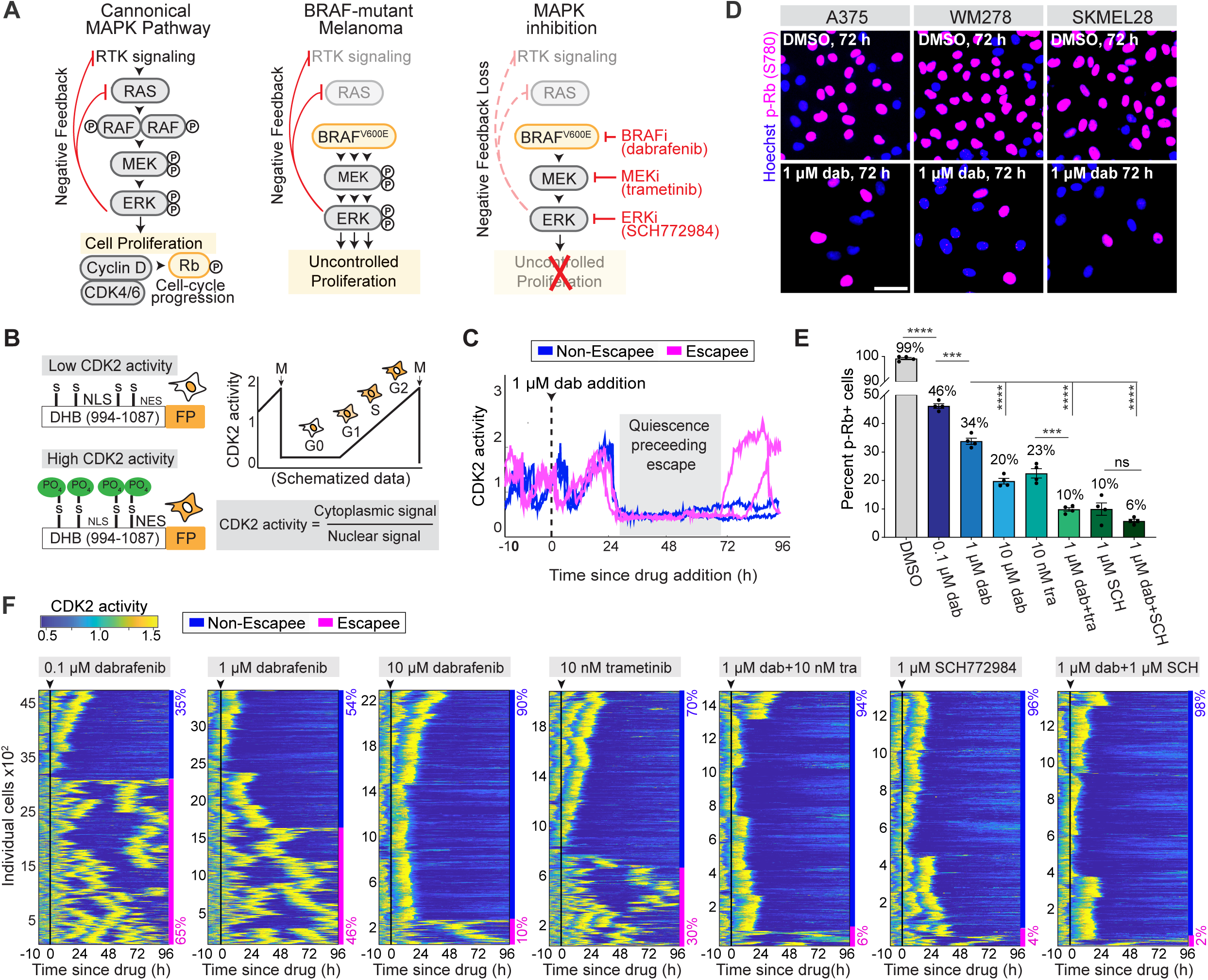
| A subpopulation of persister cells can rapidly and non-genetically escape from MAPK inhibition to resume proliferation. **(A)** Signaling diagrams linking MAPK pathway activity to cell-cycle regulation in normal (canonical) and BRAF^V600^-mutant melanoma cells +/-drug treatment. p-Rb is used to report cell-cycle commitment throughout this study. MAPK pathway inhibitors used in this study are shown. Red curves indicate negative feedback loops. **(B)** Schematic of CDK2 activity sensor for cell-cycle status (*38*). Phosphorylation of the sensor by CDK2 masks the nuclear localization signal (NLS) and unmasks the nuclear export signal (NES), causing translocation of the sensor to the cytoplasm. The cytoplasmic/nuclear ratio serves as a readout for CDK2 activity, which rises as cells move through the cell cycle and turns off when cells exit the cell cycle into quiescence. **(C)** Representative single-cell traces of CDK2 activity in 1 µM dabrafenib-treated A375 escapees (pink) and non-escapees (blue). **(D)** Representative immunofluorescence (IF) images of three human BRAF^V600E^-mutant melanoma cell lines 72 h after 1 μM dabrafenib, stained for DNA content (Hoescht) and p-Rb (S780). Scalebar = 50 μm. **(E)** Percentage of cycling (p-Rb^+^) A375 cells in the indicated drug conditions. Cells were treated for 72 h. Error bars: mean ± std of four replicate wells; *p*-values determined by unpaired t-test between the indicated drug conditions, specified with a black line linking the conditions. **(F)** Heatmaps of temporal CDK2 activities in thousands of single A375 cells, before and after treatment with the indicated drug condition. Arrow and black line mark time of drug addition. Each row represents the CDK2 activity in a single cell over time according to the colormap. The percentages mark the proportion of cells with each behavior (pink, escapee; blue, non-escapee). Sample sizes are defined on y-axes.

Due to this inherent link between MAPK signaling and cell-cycle entry, MAPK pathway mutations are drivers of uncontrolled proliferation in many cancers. In melanoma, >50% of cases are driven by a BRAF^V600^ mutation (*3*), which causes constitutive MAPK signaling, independent of upstream RTK activation (Fig. 1A, *middle*). Clinically, targeted inhibitors of mutant BRAF^V600^ (dabrafenib, encorafenib) (*4*, *5*) and MEK (trametinib, binimetinib) (*6*) are the primary treatment for melanoma patients who are refractory to standard-of-care immunotherapy (*7*, *8*) (Fig. 1A, *right*). While initially effective, BRAF/MEK therapies inevitably lead to drug resistance and tumor relapse. Emerging studies increasingly link this drug resistance to residual drug-tolerant persister cells that survive the initial treatment (*9–12*). While some cells may survive due to pre-existing genetic mutations or epigenetic cell states (intrinsic drug resistance) (*13–15*), others can non-genetically adapt *in response* to drug pressure by rewiring their internal signaling cascades to tolerate drug (acquired drug resistance). (*16–20*).

As a result of these transient and heterogeneous features, identifying and studying drug-tolerant cells is extremely challenging. However, using live and fixed-cell imaging, we and others have tracked the behavior of drug-treated BRAF^V600E^ melanoma cells (*14*, *17*, *21–23*). Using a live-cell sensor for CDK2 activity to read out proliferation-quiescence decisions in living single cells (Fig. 1B), we found significant heterogeneity *within* persister populations. While all persisters survive drug and enter a quiescent state, a subset of cells termed “escapees” can escape drug action to resume proliferation within 3-4 days (*21*) (Fig. 1C). This distinction between escapees and non-escapees is further complicated by significant heterogeneity within the escapee population in terms of their timing of cell-cycle re-entry. Importantly, escapees revert to parental drug sensitivity upon drug withdrawal, indicating that their behavior is driven by plasticity in response to drug pressure, not pre-existing genetic mutations (*21*, *24*). Therefore, while escapees and non-escapees appear indistinguishable during the drug-induced quiescent state, heterogeneity in the timing and propensity for escape among these cells likely reflects underlying differences in cell signaling that precede and dictate the emergence of these distinct fates.

As a result, it is crucial to identify the signaling events that precede a persister cell’s commitment to distinct cell-cycle outcomes, as early dynamics may reveal critical determinants underlying these specific fates. While single-cell transcriptomic and bulk proteomic techniques have powerfully contributed to understanding features of drug escape, these approaches have several limitations. First, these methods only capture a static snapshot of signaling activity, whereas signaling is dynamic and integrated over time to guide proliferation-quiescence decisions (*25–27*). For example, signaling from RTKs to ERK occurs on the timescale of minutes, yet cell-cycle re-entry occurs only every 10+ hours. Thus, the cumulative and integrated dynamics of ERK signaling better inform a cell’s fate than a single snapshot of ERK activity (*28*, *29*). Second, these techniques cannot track the temporal progression of fate decisions. Without this temporal context, linking signaling activity in quiescent cells to their future fate is difficult, as cells’ fates are unknown during the quiescent period while the drug is still effective. Additionally, when cells are analyzed after fate divergence (e.g., after escapees and non-escapees emerge), it is unclear whether observed signaling differences are true causes of escape or merely consequences of cell-cycle status.

To overcome these limitations, we employed time-lapse imaging combined with an automated single-cell tracking pipeline, EllipTrack (*30*), to monitor drug-treated melanoma cells co-expressing reporters for real-time signaling dynamics and a CDK2 activity sensor for cell-cycle status (Fig. 1B). This approach enabled us to link dynamic signaling events in drug-induced quiescent cells to future cell-cycle fate decisions. Unexpectedly, we found only a tenuous connection between integrated ERK activity and cell-cycle re-entry, suggesting that MAPK activity is necessary but insufficient for driving escape. Instead, we identified mTORC1 signaling as an additional essential mediator of escape, enhancing protein translation and growth rates during the quiescent period of future escapees. By leveraging the heterogeneity in escape timing, we discovered that persisters exhibit varying proliferative potential, or cell-cycle re-entry “inertia”, reflective of their ability to reactivate both ERK and mTORC1 signaling, which converge to drive the accumulation of Cyclin D1 protein, a key bottleneck for cell-cycle re-entry. Taken together, our data highlight how the integration of multiple complex signaling pathways drives specific cell-cycle fate decisions in drug-treated melanoma cells.

## RESULTS

### A subpopulation of cells can rapidly and non-genetically escape from MAPK inhibition to resume proliferation

To first assess the prevalence of cell-cycle heterogeneity across melanoma lines, we profiled three BRAF^V600E^ mutant melanoma lines (A375, WM278, and SKMEL28) (Fig. S1A) before and after treatment with dabrafenib, a potent BRAF^V600^ inhibitor. Upon immunostaining for hyperphosphorylated Rb (p-Rb), a marker of cell-cycle commitment (*31*), we found that all cell lines contained a subpopulation of proliferative escapees after 72 hours of drug treatment (Fig. 1D). To assess the possibility that this effect resulted from suboptimal inhibition of the MAPK pathway, we evaluated all three cell lines across increasing concentrations of dabrafenib, as well as the clinically relevant combinations of dabrafenib with either a MEK (trametinib) or an ERK (SCH772984) inhibitor (*32*). Consistent with previous findings, co-inhibition of BRAF and MEK or ERK reduced the fraction of escapees (Fig. 1E-F; Fig. S1B-D), highlighting the MAPK pathway’s key role in driving cell proliferation. However, despite this combination therapy or treatment with supraphysiological concentrations of a single drug, a small subpopulation of escapees was still observed, underscoring a critical need to understand how these cells are rewiring their signaling pathways to resume proliferation in the presence of these drugs.

### MAPK pathway reactivation is necessary but not sufficient for driving cell-cycle re-entry in the presence of drug

Extensive literature has linked melanoma cell-cycle re-entry in the presence of BRAF inhibitors to “paradoxical MAPK reactivation,” where drug-induced suppression of MAPK signaling relieves negative feedback on upstream RTKs/Ras, priming them for reactivation (*15*, *18*, *23*, *33–35*). This upstream Ras reactivation triggers the dimerization of BRAF^V600^ monomers with wild type RAF monomers (*36*). Since first-generation BRAF inhibitors like dabrafenib only inhibit BRAF^V600^ monomers, this dimerization enables drug evasion (Fig. 2A). To investigate whether MAPK reactivation underlies the escapee phenotype, we immunostained drug-treated A375 cells for p-ERK (T202/Y204) to assess ERK activity and p-Rb (S780) as a marker for cell-cycle commitment. Single-cell quantification of ERK phosphorylation showed complete suppression of ERK activity within 2 hours of dabrafenib or dabrafenib/trametinib, followed by a partial rebound over the next 72 hours (Fig. 2B-C). Under dabrafenib, this rebound was accompanied by a long-tailed distribution of p-ERK, with some cells reactivating ERK to drug-naïve levels (Fig. 2C, *left*), likely in a pulsatile fashion (*23*, *37*). Indeed, combination dabrafenib/trametinib suppressed these ERK-reactivating cells, although not completely (Fig. 2C, *right*). Notably, the rebound in p-ERK temporally coincided with the emergence of escapees (p-Rb^+^ cells) (Fig. 2B), supporting the notion that paradoxical MAPK reactivation drives escape under drug treatment. However, when we compared p-ERK levels between escapees (p-Rb^+^) and non-escapees (p-Rb^-^), significant overlap in staining distributions was observed in the A375, WM278, and SKMEL28 cell lines (Fig. 2D-E). These data show that p-ERK levels are not generally higher in p-Rb^+^ escapees compared to p-Rb^-^ non-escapees.

**Figure. 2.**
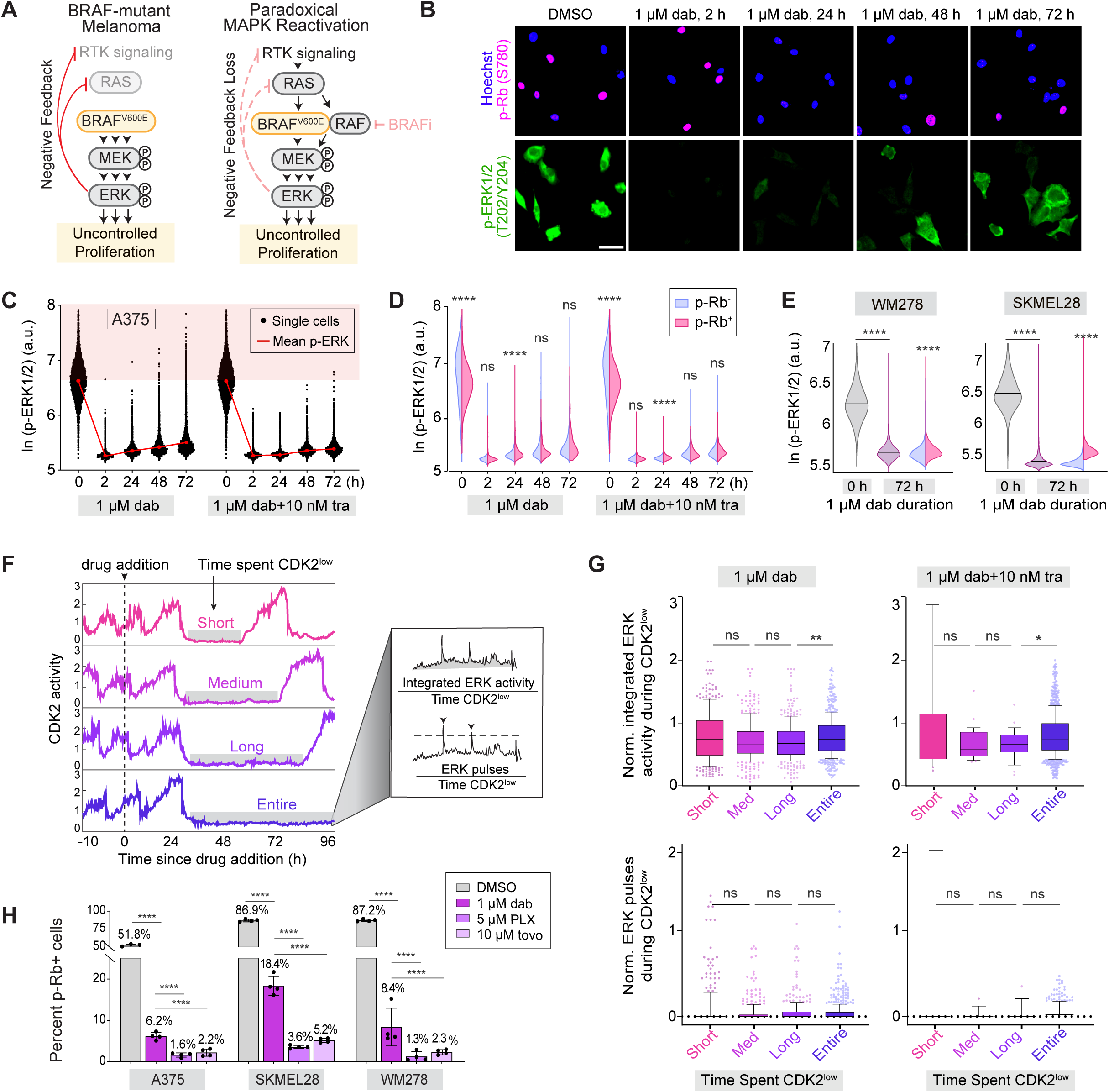
| MAPK pathway reactivation is necessary but not sufficient for driving cell-cycle re-entry in the presence of drug. **(A)** Signaling diagrams linking negative feedback (red) to MAPK pathway reactivation. **(B)** Representative IF images of A375 cells co-stained for Hoechst, p-Rb (S780), and p-ERK1/2 (T202/Y204) after 1 µM dabrafenib for 0, 2, 24, 48, or 72 h. Scalebar = 50 μm. **(C)** Single-cell violin plots of p-ERK1/2 intensity over time after 1 µM dabrafenib or combination 1 µM dabrafenib + 10 nM trametinib in A375 cells. Red shading is drawn at the mean intensity for untreated cells. Red line connects the mean intensity at each time point. n > 7,000 cells per violin. **(D)** Re-analysis of bulk violins in Fig. 2C as split violins to determine p-ERK1/2 intensity in non-escapees (p-Rb^-^, blue) vs escapees (p-Rb^+^, pink). n > 185 cells per violin. **(E)** Bulk violins of p-ERK1/2 intensity 72 h after no treatment (gray) or 1 μM dabrafenib (purple) in WM278 and SKMEL28 cells. n > 5,000 cells per violin. Split violins separate p-ERK1/2 intensity in non-escapees (p-Rb^-^, blue) vs escapees (p-Rb^+^, pink). n >300 cells per violin. **(F)** Schematic representing how MAPK activity will be linked to cell-cycle re-entry timing. Heterogeneity in 1 μM dabrafenib-treated A375 cells regarding time spent CDK2^low^ (>0.8 cutoff) is visualized in representative CDK2 activity traces. To quantify MAPK activity during the CDK2^low^ period, the integrated ERK activity (area under the trace) and number of ERK pulses were measured via the ERK KTR_C/N_ and normalized for time spent CDK2^low^. **(G)** Box plots of time-normalized ERK activity and number of ERK pulses during the CDK2^low^ period of the indicated drug conditions in A375 cells. Escapees were clustered according to time spent CDK2^low^ (short, medium, or long), with each group representing the bottom, middle, and top 20% of CDK2^low^ duration lengths. n = 352 cells (1 μM dabrafenib) and n = 29 cells (1 μM dabrafenib + 10 nM trametinib). Non-escapees, which remained CDK2^low^after drug treatment for the remainder of the movie, were grouped together. n = 645 (1 μM dabrafenib) and n >1100 cells (1 μM dabrafenib + 10 nM trametinib). Box displays median and quartiles, whiskers display 10-90^th^ percentile. **(H)** Percentage of escapees (p-Rb^+^) cells in the indicated drug conditions after 72 h. Error bars: mean ± std of four replicate wells. For bar graphs, *p*-values were calculated by unpaired t-tests between indicated drug conditions, specified with a black line linking the conditions. For bulk violins, *p*-values were determined by Mann-Whitney U-tests between indicated drug conditions, specified with a black line linking the conditions. For split violins, *p*-values were determined by Mann-Whitney U-tests between non-escapee (p-Rb^-^) compared to escapee cells (p-Rb^+^).

This observation is explainable, however, since ERK activity at a snapshot in time carries little information about cell-cycle status – pRb+ cells can be high or low for p-ERK, as can pRb-cells (Fig. S2A). Previous studies have suggested that it is the integrated ERK activity or ERK pulses over time that dictate proliferation-quiescence decisions (*25*, *37*). Despite several attempts to make this connection in drug-treated cancer cells, no study has quantitatively measured and compared the integrated ERK activity/pulses in the drug-induced quiescence period preceding the emergence of escapees and non-escapees.

To achieve this goal, we created A375 cells stably expressing an H2B-mIFP nuclear marker for cell tracking, DHB-mCherry (CDK2 activity sensor) for cell-cycle status (*38*, *39*) (Fig. 1B), and Elk1-mClover (ERK KTR) to measure ERK activity (*40*). When phosphorylated, the ERK KTR sensor translocates from the nucleus to the cytosol, providing a time-resolved, single-cell readout of kinase activity. Importantly, we recently identified crosstalk from CDK2 onto the ERK KTR sensor, confounding ERK activity measurements (*41*). However, because CDK2 activity only rises once cells commit to the cell cycle (*38*), this crosstalk only affects cycling cells, allowing the ERK KTR sensor to accurately measure ERK activity in quiescent cells prior to cell-cycle re-entry. We therefore performed time-lapse imaging on the 3-color A375 cells treated with either dabrafenib or the dabrafenib/trametinib combination. We quantified the integrated ERK activity (area under the trace) and the number of ERK pulses during the quiescence preceding cell-cycle re-entry and normalized these values to the time spent CDK2^low^ for each cell (Fig. 2F). While dabrafenib/trametinib treatment reduced mean ERK activity (Fig. S2B), the number of ERK pulses (Fig. S2C, *right*), and the fraction of escapees (Fig. S2C, *left*), compared to dabrafenib alone, we found no correlation between time-normalized integrated ERK activity (Fig. 2G, *top*) or time-normalized number of ERK pulses (Fig. 2G, *bottom*) and the time cells spent CDK2^low^ in either drug condition. These data suggest that while ERK is indeed getting paradoxically reactivated in the presence of drug as a mechanism to aid escape, it is getting reactivated to similar levels in non-escapees and future escapees during the quiescent period.

This finding led us to hypothesize that ERK activity is necessary but not sufficient for driving cell-cycle re-entry in the presence of drug. Rather, future escapee cells may be upregulating other signaling pathways to compensate for reduced MAPK activity and to drive proliferation, relative to non-escapees. Further supporting this idea, treatment with the paradox breaker PLX8394 (*42*) and the pan-RAF inhibitor tovorafenib (*43*), both designed to prevent paradoxical MAPK reactivation, still resulted in subpopulations of escapee cells capable of evading drug and resuming proliferation. (Fig. 2H; Fig. S1B-D).

### Escapees are enriched for markers of mTORC1 activity relative to non-escapees

To elucidate potential pathways additionally enabling drug escape, we previously performed single-cell RNA sequencing (scRNA-seq) in A375 cells treated with dabrafenib for 72 hours (*21*). Pathway enrichment analysis using the Hallmark Gene Set identified mTORC1 (mechanistic target of rapamycin complex 1), involved in cell growth and protein translation (*44*), as the top pathway unique to escapees compared to non-escapees and untreated cycling cells (*21*) (Fig. 3A). Notably, the scRNA-seq analysis also identified pathways previously linked to persister cells including the epithelial-to-mesenchymal transition pathway (*45*, *46*), p53 activation (*47*), and oxidative stress (*48*, *49*), among others. Important in the context of drug treatment, the mTORC1 pathway has been shown to sense microenvironment changes and accordingly adjust the balance between cellular anabolic (protein, lipid, nucleotide production) and catabolic (TFEB-mediated autophagy suppression) processes to regulate cell growth and proliferation (*50–53*) (Fig. 3B).

**Figure. 3.**
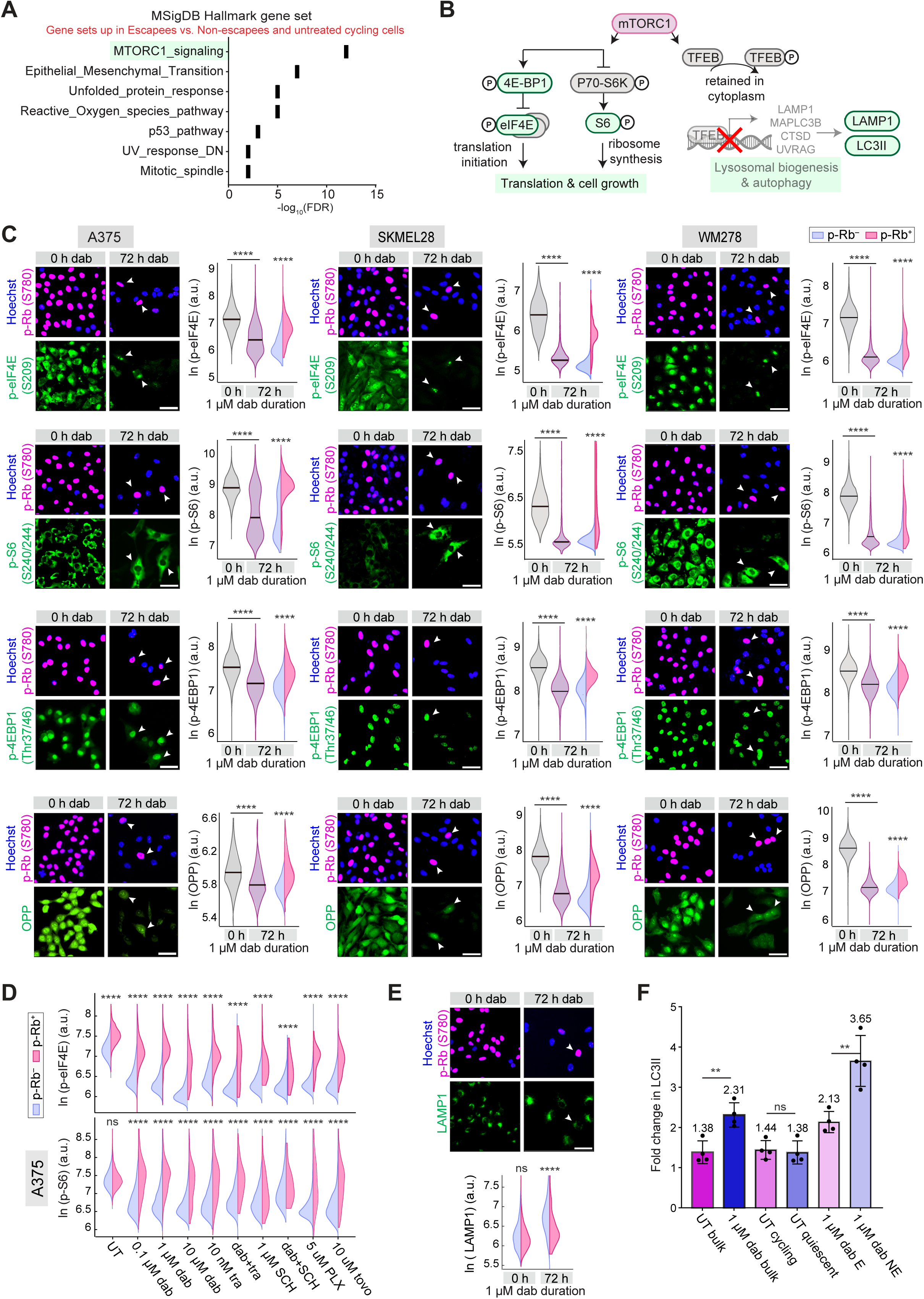
| Escapees are strongly enriched for markers of mTORC1 activity. **(A)** MSigDB hallmark gene set enrichment analysis of 40 differentially expressed genes from scRNA-sequencing dataset comparing 1 µM dabrafenib-treated escapees to non-escapees and untreated cycling cells. A false-discovery rate cutoff of 0.05 was used. Data replotted from (*21*) **(B)** Signaling diagram of mTORC1 pathway on growth and translation vs. autophagy. **(C)** Representative IF images of either A375, SKMEL28, or WM278 cells treated with 1 µM dabrafenib for 72 h and co-stained for Hoescht, p-Rb (S780), and various mTORC1 activity markers (p-eiF4e (S209), p-S6 (S240/244), p-4EBP1 (Thr37/46)), and translation rate (using OPP). Quantification of each marker is plotted as bulk violin plots in untreated cells (gray) vs dabrafenib-treated (purple) cells, as well as split violin plots in dabrafenib-treated non-escapees (p-Rb^-^, blue) vs. escapees (p-Rb^+^, pink). Scalebar = 50 μm. White arrows highlight representative escapee cells. **(D)** Split violin plots quantifying two mTORC1 activity markers in non-cycling (p-Rb^-^, blue) vs cycling A375 cells (p-Rb^+^, pink) treated for 72 h with varying drug conditions. UT, untreated. **(E)** Representative IF images and split violin plots of untreated and 1 µM dabrafenib-treated A375 cells quantifying lysosomal marker LAMP1 in non-escapees (p-Rb^-^, blue) vs escapees (p-Rb^+^, pink). Scalebar = 50 μm. **(F)** Autophagic flux measured in UT and 1 µM dabrafenib-treated A375 by change in LC3II after a 3 h treatment with 10 μM chloroquine. Error bars: mean ± std of four replicate wells. For bar graphs, *p*-values were calculated by unpaired t-tests between indicated drug conditions, specified with a black line linking the conditions. For bulk violins, *p*-values were determined by Mann-Whitney U-tests between indicated drug conditions, specified with a black line linking the conditions. For split violins, *p*-values were determined by Mann-Whitney U-tests between non-escapee (p-Rb^-^) compared to escapee cells (p-Rb^+^).

To assess a possible role for mTORC1 in mediating drug escape, we profiled three BRAF^V600E^ mutant cell lines (A375, SKMEL28, WM278) with and without dabrafenib for 72 hours and performed multiplexed immunofluorescence imaging for the proliferation marker p-Rb, markers of increased mTORC1 activity (p-S6, p-eIF4E, p-4E-BP1) (*54–57*), and the rate of global protein translation using the O-propargyl puromycin (OPP) assay (Fig. 3C). As expected, markers of mTORC1 activity and translation rate were reduced at the bulk population level in dabrafenib-compared to DMSO-treated cells, highlighting the limitation of previous methods in identifying markers of escape in drug-treated populations where cycling cells represent only a small fraction of the population. However, when we separated persisters by cell-cycle status, we found that escapees (p-Rb^+^) were significantly enriched for every mTORC1 marker, as well as for protein translation rate (OPP), compared to non-escapees (p-Rb^-^) (Fig. 3C). We additionally observed the same results when cells were treated for 72 hr with dabrafenib, trametinib, the ERK inhibitor SCH772984, the paradox breaker PLX83954, or the pan-RAF inhibitor tovorafenib, alone or in combination (Fig. 3D).

Importantly, mTORC1 promotes cell growth not only by stimulating biosynthetic pathways, but also by inhibiting cellular catabolism through repression of the lysosomal-mediated autophagic pathway. Indeed, mTORC1 directly regulates TFEB, the master transcription factor for lysosome biogenesis, via an inhibitory phosphorylation that sequesters TFEB to the cytoplasm, reducing TFEB-mediated transcription of lysosomal and autophagy related genes (*51*, *58*) (Fig. 3B). To test for differences in autophagic flux between escapee and non-escapee persisters, we fixed and stained 72-hour dabrafenib-treated A375 cells for p-Rb and either the lysosomal marker LAMP1 or LC3II (*59*) with and without a 3-hour treatment with 10 μM of the autophagy inhibitor chloroquine. We identified a marked decrease in LAMP1 and autophagic flux in cycling escapees compared to non-cycling non-escapees, which we reasoned was likely due to increased mTORC1-mediated cytoplasmic suppression of TFEB, and thus, decreased transcription of lysosomal and autophagy-related genes in escapees compared to non-escapees (Fig. 3E-F).

### Growth Rate during the CDK2^low^ period is correlated with drug escape both within and across drug conditions

Previous work has highlighted the importance of cell growth and biomass thresholds in driving cell-cycle commitment in yeast and in normal human cells (*60–64*). Our results therefore led us to hypothesize that the extent of mTORC1 activity controls cellular anabolic and catabolic activity, and thus cell mass and cell growth, potentially underlying the observed differences in persister cell-cycle fate. To test this hypothesis, we measured differences in biomass accumulation, also known as the growth rate, during the CDK2^low^ period preceding cell-cycle re-entry in drug-treated melanoma cells. Previous work has shown that changes in the dry biomass of a cell directly correlate to changes in the nuclear volume and nuclear area of a cell over time, providing a functional proxy for growth rate in live-cell imaging experiments (*60*, *65–67*). Consistently, DMSO and 72-hour dabrafenib-treated A375 cells stained with AlexaFluor 488-Succinimidyl Ester (SE-A488), a quantitative protein stain for cell mass (*60*, *65*), showed a direct correlation between cell mass (SE-A488 levels), cytoplasmic volume, nuclear volume, and nuclear area (Fig. S3A). Conveniently, the change in nuclear area over time as a readout for growth rate is freely obtained in our live-cell imaging experiments by segmenting and tracking our nuclear marker, H2B-mIFP.

Using this approach, we re-analyzed the live-cell, time-lapse imaging movies of A375 cells treated with multiple MAPK pathway inhibitors (Fig. 1F) and quantified the growth rate (change in nuclear area) specifically during the drug-induced CDK2^low^ period preceding escape (Fig. S3B). We found that the growth rate during this quiescence period directly correlated with the timing of cell-cycle re-entry, where faster growing quiescent cells escaped from drug to re-enter the cell cycle earlier than slower growing quiescent cells (Fig. 4A-B). To confirm that these differences in nuclear area and cell-cycle re-entry time reflected true biological differences in persister cell growth rates, we tested several targeted perturbations known to directly alter growth rate. We performed time-lapse imaging of H2B-mIFP and DHB-mCherry expressing A375 cells treated with either DMSO, dabrafenib, an mTORC1 inhibitor RapaLink-1 (*68*), or the combination (Fig. 4C). As mTORC1 directly upregulates pathways involved in biomass synthesis and cell growth, inhibition of mTORC1 via RapaLink-1 directly reduces cell growth, which was captured by our growth rate measurements across all RapaLink-1 treated conditions (Fig. 4D). We additionally found that both single agent and combination RapaLink-1 plus dabrafenib reduce the fraction of escapees in A375, SKMEL28, and WM278 cells (Fig. 4C; Fig. S3C), suggesting a role for mTORC1 in driving drug escape. Indeed, we identified a strong dose-dependent relationship between the average cellular growth rate during the CDK2^low^ period and the percentage of cycling cells across increasing levels of MAPK pathway inhibition (Fig. 4E).

**Fig. 4.**
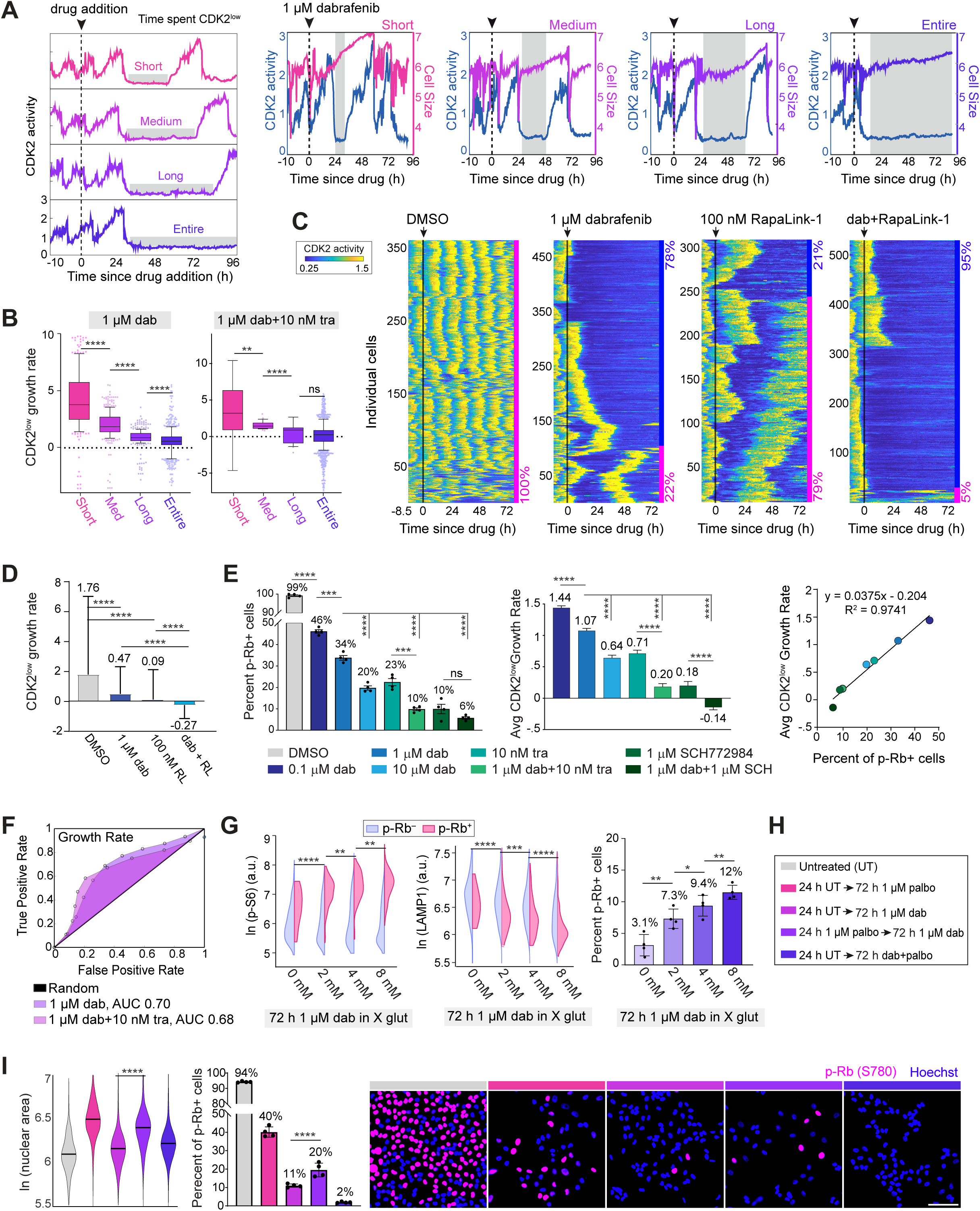
| Growth rate during the CDK2^low^ period is correlated with drug escape both within and across drug conditions. **(A)** Representative single cell traces of temporal CDK2 activities in 1 μM dabrafenib-treated A375 cells overlaid with corresponding growth rates (change in nuclear area over time), during the CDK2^low^ period (gray boxes) preceding escape. **(B)** Re-analysis of data from Fig. 2G as box plots displaying growth rates during the CDK2^low^ period of 1 μM dabrafenib-treated or 1 µM dabrafenib + 10 nM trametinib-treated A375 cells. Clustering and sample size is retained from Fig. 2G. **(C)** Heatmaps of temporal CDK2 activities in hundreds of single A375 cells, before and after treatment with the indicated drug conditions. Arrow and black line mark time of drug addition. Each row represents the CDK2 activity in a single cell over time according to the colormap. The percentages mark the proportion of cells with each behavior (pink, escapee; blue, non-escapee). Sample sizes are defined on y-axes. **(D**) Quantification of the average growth rate for each indicated drug condition (average of each cell’s growth rate across its CDK2^low^ period) from time-lapse imaging in Fig. 4C. **(E)** Correlation between the percentage of escapees (p-Rb^+^) after 72 h (reproduced from Fig. 1E) vs. the average growth rates during the CDK2^low^ period across the indicated drug conditions in A375 cells. Error bars: mean ± std of four replicate wells. R value is calculated as the Pearson correlation coefficient. **(F)** ROC analysis using growth rates from Fig. 2G to predict future non-escapee and escapee cells in 1 μM dabrafenib-treated or 1 µM dabrafenib + 10 nM trametinib-treated A375 cells. AUC indicates the area under the curve for each condition. **(G)** A375 cells treated with 1 μM dabrafenib for 72 h in increasing concentrations of glutamine (standard concentration is 4 mM) and stained for p-S6 (S240/244), LAMP1, and p-Rb (S780). For split violins, *p*-values were determined by Mann-Whitney U-tests between escapees (p-Rb^+^) from the indicated drug conditions, specified with a black line. **(H)** Schematic of experimental drug conditions with associated legend. **(I) Left:** Quantification of nuclear area (bulk violins) and percentage of escapees (p-Rb^+^) (bar graphs) in the indicated drug conditions. Error bars: mean ± std of four replicate wells. **Right:** Representative immunofluorescence images of A375 cells after the indicated drug conditions, stained for Hoescht and p-Rb (S780). Scale bar = 100 μm. For bar graphs, *p*-values were calculated by unpaired t-tests between indicated drug conditions, specified with a black line linking the conditions. Unless otherwise specified, for bulk and split violins, *p*-values were determined by Mann-Whitney U-tests between indicated drug conditions, specified with a black line linking the conditions.

To evaluate whether the growth rate prior to cell-cycle re-entry can predict drug escape, we measured the growth rate of each cell during its entire CDK2^low^ period (Fig. 4A-B) to construct receiver operating characteristic (ROC) curves, comparing true-positives (escapees) against false-positives (non-escapees) across varying detection thresholds (e.g. growth rates) (Fig. 4F). This analysis revealed that growth rate was moderately predictive of escape, with an AUC of 0.70 in dabrafenib-treated cells and 0.68 in combination dabrafenib plus trametinib-treated cells (Fig. 4F). While these results suggest that growth rate has moderate resolving power for distinguishing escapees from non-escapees, this may underestimate its true predictive potential. This could occur because performing a binary classification of cells into escapees and non-escapees (rather than distinguishing early, mid, and late escapees) fails to account for the fact that late-escaping cells often exhibit growth rates similar to non-escapees (Fig. 4A-B), limiting the resolving power of growth rate on predicting escape.

While these previous experiments clearly highlight the correlative and predictive role of mTORC1 activity and growth rate on drug escape, we wanted to directly test their causal involvement. To establish a causal role for mTORC1 in driving escape, we treated A375 cells with DMSO (Fig. S3D) or dabrafenib (Fig. 4G) for 72 hours in the presence of increasing concentrations of glutamine, a known activator of mTORC1 (*69*). Immunostaining revealed that under drug treatment, higher glutamine levels increased mTORC1 activity, as indicated by elevated p-S6 staining and reduced LAMP1 staining, which was not observed in DMSO-treated cells. These changes were additionally accompanied by a higher percentage of cycling persisters, correlating elevated mTORC1 activity with drug escape (Fig. 4G; Fig. S3D).

Building on these findings, we hypothesized that mTORC1 drives escape by enhancing cell growth and protein translation. To test if increased cellular growth is causally linked to cell-cycle re-entry in the presence of drug, we increased cell mass by pre-treating cells for 24 hours with the CDK4/6 inhibitor palbociclib (*70*), as done previously (*25*). We then washed off the palbociclib and treated cells for 72 hours with dabrafenib (Fig. 4H; Fig. S3E). Palbociclib promotes cell growth by stalling cells in G0 to accumulate protein mass in the absence of cell-cycle progression (*71*, *72*), as evidenced by the observed increase in nuclear area following pre-treatment (Fig. 4I, *left*). We reasoned that palbociclib pre-treated cells would effectively have a “head start” in accumulating mass, and thus would be primed to escape dabrafenib-induced quiescence more readily. Indeed, we found that the fraction of escapees increased 2-fold when cells were pre-treated with palbociclib (Fig. 4I) indicating that ERK activity, mTORC1-mediated cell growth and translation, and cell-cycle commitment are inter-dependent signals that determine cell-cycle fate upon MAPK inhibition.

### Cyclin D1 expression is a key bottleneck for cell-cycle re-entry in the presence of MAPK pathway inhibitors

While our data clearly indicate that both the MAPK and mTORC1 pathways contribute to drug escape, it remains unclear how melanoma cells integrate these complex signaling pathways under prolonged drug treatment to make proliferation-quiescence decisions. To determine this, we aimed to identify a bottleneck where these pathways converge to regulate cell-cycle re-entry. Previous work from our lab has shown that ERK signaling dynamics are integrated in non-cancerous cells through the rate of Cyclin D1 protein production, which determines the probability of cell-cycle entry (*25*). This led us to question whether the accumulation of Cyclin D1, influenced by both MAPK and mTORC1 activity, may similarly represent a key bottleneck for cell-cycle re-entry in drug-treated melanoma cells.

Importantly, Cyclin D1 sits at the nexus between the MAPK and mTORC1 signaling pathways and cell-cycle re-entry. Specifically, Cyclin D1 is transcriptionally regulated downstream of ERK activity and translationally regulated downstream of mTORC1 activity (*73–77*) (Fig. 5A). In support of this, treatment of A375 cells with increasing doses of dabrafenib resulted in greater suppression of Cyclin D1 mRNA, and consequently, Cyclin D1 protein levels. In contrast, increasing doses of the mTORC1 inhibitor RapaLink-1 had no effect on Cyclin D1 mRNA levels, but significantly reduced Cyclin D1 protein levels (Fig. 5B). This regulation is critical, as Cyclin D1’s short half-life (∼30 minutes) (*25*, *78*) renders it highly sensitive to fluctuations in protein synthesis rates, allowing rapid adjustments to signaling events and highlighting its role as a key regulatory node in driving cell-cycle entry. Indeed, we showed that Cyclin D1 plays a causal role in cell-cycle commitment: its knockdown reduced, while its overexpression increased, the percentage of cycling cells in both DMSO– and dabrafenib-treated A375 cells (Fig 5C; Fig. S4A). When we further suppressed the MAPK pathway, the abundance of Cyclin D1 protein within the subpopulation positioned to escape (i.e. the currently quiescent p-Rb^-^ cells) was further reduced in a dose-dependent fashion. Thus, low levels of Cyclin D1 protein likely help maintain cell-cycle withdrawal in non-escapees (Fig. 5D-E).

**Fig. 5.**
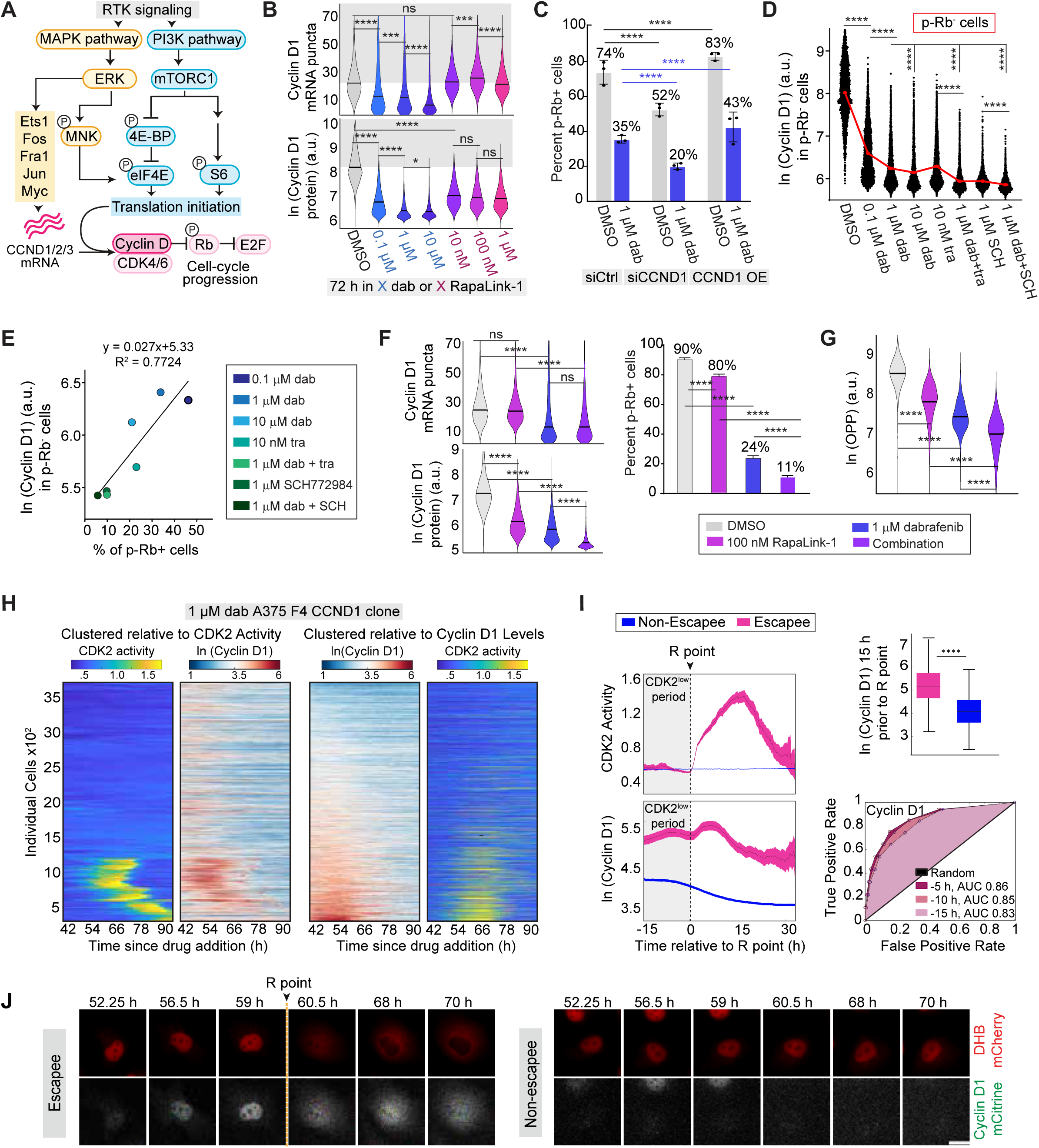
| Cyclin D1 expression is a key bottleneck for cell-cycle re-entry in the presence of MAPK pathway inhibitors. **(A)** Signaling diagram for Cyclin D1 regulation from the MAPK pathway (yellow) and the mTORC1 pathway (blue). **(B)** Bulk violins of Cyclin D1 mRNA (top, quantified by RNA-FISH) and Cyclin D1 protein (bottom, quantified by immunofluorescence) in A375 cells treated for 72 h with either DMSO or increasing concentrations of dabrafenib (blue shades) or RapaLink-1 (pink shades). **(C)** Percentage of cycling A375 cells (p-Rb^+^) quantified 72 h after treatment with either DMSO or 1 μM dabrafenib and 24 h after transfection with either siControl, siCCND1 knockdown, or Cyclin D1 overexpression constructs. Specifically, transfection mixture +/-dabrafenib was added to cells after 48 h in dabrafenib or DMSO and removed 6 h later. Cells were then cultured in dabrafenib or DMSO for another 18 hours. **(D)** Cyclin D1 protein levels in non-cycling (p-Rb^-^ cells), quantified by immunostaining of Cyclin D1 and p-Rb (S780) after 72 h in the indicated drug conditions. **(E)** Correlation between the percentage of cycling A375 cells (p-Rb^+^) and the average Cyclin D1 protein levels in non-cycling cells (p-Rb^-^) after 72 h in the indicated drug conditions. Error bars: mean ± std of four replicate wells. R value is calculated as the Pearson correlation coefficient. **(F)** Cyclin D1 mRNA levels (left top), Cyclin D1 protein levels (left bottom), percentage of cycling cells (p-Rb^+^) (right), and **(G)** protein translation rate by OPP assay in A375 cells after 72 h in the indicated drug conditions. **(H)** Heatmaps of temporal CDK2 activities (yellow/blue heatmaps) and endogenous mCitrine-Cyclin D1 (red/blue heatmaps) in thousands of single A375 cells after 48 h in 1 μM dabrafenib. The left two heatmaps are clustered according to CDK2 activity and the right two heatmaps are clustered according to Cyclin D1 protein levels. Sample sizes are defined on y-axes. **(I) Left:** Population average and 95% confidence interval of CDK2 activity and endogenous mCitrine-Cyclin D1 in 1 μM dabrafenib-treated A375 cells aligned to the restriction point (R point, the point of CDK2 activity rise) in order to visualize endogenous Cyclin D1 levels prior to escape in escapees (pink) vs non-escapees (blue). **Right:** Cyclin D1 levels 15 h prior to escape are quantified in the boxplots (right top) in escapees vs non-escapees. Box displays median and quartiles, whiskers display 10-90^th^ percentile. ROC analysis using mCitrine-Cyclin D1 at different timepoint (–5, –10, and –15 h) prior to escapee R point to predict future non-escapee and escapee cells (right bottom). AUC indicates the area under the curve for each condition (right bottom). **(J)** Film strip from Movie S1A highlighting mCitrine-Cyclin D1 levels relative to CDK2 activity (DHB-mCherry) in one escapee (left) vs one non-escapee (right). Arrow and yellow dotted line mark restriction (R) point. For bar graphs, *p*-values were calculated by unpaired t-tests between indicated drug conditions, specified with a black line linking the conditions. For bulk violins, *p*-values were determined by Mann-Whitney U-tests between indicated drug conditions, specified with a black line linking the conditions.

These findings suggest that while MAPK inhibition reduces Cyclin D1 mRNA levels, mTORC1 activity may compensate by driving Cyclin D1 translation, enabling its accumulation over time, even under drug treatment. This raises the possibility that the previously observed reduction in cycling cells with dual MAPK and mTORC1 inhibition (Fig. 4C; Fig. S3C), is driven by reduced Cyclin D1 translation, which would indicate a critical role for mTORC1 in sustaining Cyclin D1 expression during MAPK inhibition. Notably, combining dabrafenib with the mTORC1 inhibitor RapaLink-1, significantly reduced Cyclin D1 protein levels compared to either single agent alone, with Cyclin D1 levels closely mirroring the percentage of cycling A375 cells in each condition (Fig. 5F). We observed the same results in SKMEL28 and WM278 cells (Fig. S3C; Fig. S4B). These reductions in Cyclin D1 protein and cycling cells are attributable to impaired protein synthesis, as confirmed by OPP staining for global translation rates (Fig. 5G) and corroborated by similar results observed with combination treatment of dabrafenib plus the translational inhibitor 4EGI-1, which blocks translation initiation-dependent eif4E/eif4G interactions (*79*) (Fig. S4C). Collectively, these results support our hypothesis that both MAPK and mTORC1 pathways are required to maintain Cyclin D1 protein levels, with MAPK inhibition impairing mRNA production (and indirectly protein production), and mTORC1 inhibition directly impairing translation of Cyclin D1 protein, emphasizing the importance of translational control in cell-cycle re-entry under MAPK inhibiting conditions.

If Cyclin D1 expression represents a key bottleneck for cell-cycle re-entry in drug, we would predict higher Cyclin D1 protein levels in the drug-induced CDK2^low^ period in escapees compared to non-escapees. To develop a system to measure endogenous Cyclin D1 protein, we used CRISPR-based genome editing to tag Cyclin D1 at its genomic locus with the mCitrine fluorescent protein in A375 cells and confirmed that we had one wild-type and one tagged allele by PCR, genomic sequencing, western blotting, and immunofluorescence (Fig. S4D-F; Data file S1). We co-expressed H2B-mIFP for cell tracking and DHB-mCherry to monitor the cell cycle. Notably, time-lapse imaging and single-cell tracking of untreated 3-color cells showed that Cyclin D1 dynamics followed a U-shape (Fig. S5A), with levels falling in G1 upon cell-cycle commitment, remaining low in S phase, and rising in G2, as observed previously in non-cancerous cell types (*25*, *78*, *80*). To minimize photobleaching (Fig. S5B) of the dim endogenous Cyclin D1 signal, A375 cells were imaged at 20X after 48 hours in dabrafenib and computationally clustered based on either CDK2 activity (Fig. 5H, left two panels) or Cyclin D1 levels (Fig. 5H, right two panels). While all dabrafenib-treated cells showed a strong decline in Cyclin D1 (Fig. S5C), clustering revealed a strong association between elevated Cyclin D1 levels and subsequent drug escape. To further test this relationship, drug-treated cells were computationally aligned to their first escape from quiescence, marked by a rise in CDK2 activity upon passage through the restriction point (*31*, *38*) (Fig. 5, I-J; Fig. S5D). Future escapee cells showed a rise in Cyclin D1 during the quiescence period preceding cell-cycle re-entry, suggesting that Cyclin D1 levels may predict escape from drug treatment. To quantitatively assess the predictive power of Cyclin D1 levels for escape, we performed an ROC analysis using Cyclin D1 levels at various time points prior to the R point to classify cells as escapees or non-escapees (Fig. 5I, *bottom right*). This analysis showed that Cyclin D1 levels up to 15 hours before cell-cycle re-entry at the R point were highly predictive of escape, with an AUC of 0.83. Thus, we conclude that Cyclin D1 represents the earliest known indication of a persister cell’s intent to escape MAPK inhibition.

## DISCUSSION

Preclinical and clinical studies indicate that melanoma progression to a fully drug-resistant state is preceded by a reversible drug-tolerant phase, which may contribute to the development of acquired resistance. Indeed, emerging work from our lab and others highlights how a subset of drug-tolerant persister cells, termed “escapees,” can non-genetically rewire their signaling pathways to escape drug-induced quiescence and resume proliferation. Despite accumulating DNA damage from cycling under drug pressure, escapees out-proliferate non-escapees over extended drug treatments, potentially serving as a reservoir for the acquisition of *bona fide* drug resistance mutations (*21*, *22*). Thus, identifying early signaling events that enable cells to escape quiescence and re-enter the cell cycle is crucial for developing strategies to delay drug resistance and prevent tumor recurrence. (*81*).

Here, we utilized computational approaches to reconstruct single-cell lineages from time-lapse imaging data, linking critical signaling pathways to cell-cycle fate decisions. Contrary to prevailing beliefs, we discovered that MAPK reactivation is necessary but not sufficient for driving cell-cycle re-entry under MAPK inhibition. Instead, mTORC1 activity, which promotes growth and protein translation, is also required. These two pathways converge to determine the proliferative inertia of persister cells, with Cyclin D1 accumulation acting as a key bottleneck for cell-cycle re-entry, consistent with prior studies in non-cancerous cells (*63*, *82*, *83*). Our findings also implicate Cyclin D1 as a potential biomarker for distinguishing future escapees from non-escapees during the quiescent period before these distinct cell-cycle fates emerge. Indeed, quiescent cells with higher growth rates and protein translation (and thus more Cyclin D1 accumulation) exhibit greater inertia, enabling them to maintain the momentum for re-entry into the cell cycle. In contrast, cells with lower growth rates or protein translation (and less Cyclin D1) are more likely to remain quiescent, highlighting the diversity of responses within drug-treated populations.

Our findings build upon the prevailing notion that paradoxical MAPK reactivation is the primary driver of escape from BRAF inhibitor therapy. Notably, this is clinically relevant as this concept has historically informed the development of additional MAPK pathway inhibitors resistant to bleed-through signaling, including MEK and ERK inhibitors, paradox breakers, and pan-RAF inhibitors (*33*, *43*, *84*, *85*). While these inhibitors can further suppress MAPK signaling and thereby offer clinical benefit, our results underscore the importance of parallel pathways, particularly mTORC1, in sustaining Cyclin D1 protein accumulation despite MAPK inhibition (*75*, *76*, *83*, *86*). This provides a mechanistic rationale for targeting Cyclin D1, a critical bottleneck for cell-cycle re-entry, as a potentially more effective strategy for preventing persister proliferation than further inhibiting MAPK signaling. Supporting this hypothesis, preclinical in vivo and in vitro studies have demonstrated that combining CDK4/6i with BRAFi/MEKi enhances therapeutic efficacy in both BRAF– and NRAS-mutant melanoma models compared to dual MAPK inhibition alone (*86–89*). In addition, a separate study (*90*) demonstrated that treatment with CDK4/6i alongside BRAFi/MEKi significantly delayed the emergence of long-term acquired drug resistance to MAPK inhibitors. This delay was primarily attributed to the prevention of early reversible states of drug tolerance, further supporting our findings that Cyclin D1 accumulation plays a critical role in enabling cells to re-enter the cell cycle and promote resistance. The promise of this approach is additionally reflected in ongoing (NCT04720768) and completed early phase clinical trials (NCT01781572, NCT02159066, NCT02065063), which demonstrate the tolerability and potential efficacy of combining CDK4/6 inhibitors with BRAFi/MEKi in melanoma.

Notably, while our work suggests that upregulation of mTORC1 leads to sufficient Cyclin D1 levels for re-entry in persister cells, the mechanisms driving mTORC1 activation in escapees remains unclear. mTORC1 is typically activated downstream of PI3K/AKT signaling (*91*), and indeed phospho-AKT staining was elevated in escapees vs. non-escapees (Fig. S6). However, there was a large degree of overlap in the phospho-AKT distributions between escapees and non-escapees, and combination PI3K/MAPK inhibition did not reduce proliferation compared to dabrafenib alone (Fig. S6). These findings suggest that alternative mechanisms may drive mTORC1 activation in persisters, potentially bypassing traditional PI3K/AKT signaling. Indeed, both ERK and CDK4/6 has been shown to directly activate mTORC1 through phosphorylation of key regulators, such as Raptor and TSC2. (*92–94*). Similarly, AMPK, a critical sensor of cellular energy levels, can activate mTORC1 in response to changes in ATP/AMP ratios, linking metabolic stress to cell-cycle progression (*95*, *96*). Additionally, mTORC1 is highly sensitive to the availability of nutrients, particularly amino acids, which activate mTORC1 through Rag GTPases to prioritize protein synthesis (*97*). These alternative mechanisms of mTORC1 activation warrant further investigation, as they may provide insight into how cells compensate under drug pressure to resume proliferation.

While we have yet to identify any pre-drug differences in future escapees vs. non-escapees in the melanoma contexts we have tested, some evidence in the literature points to intrinsic non-genetic differences in cell state or molecular context at the time of drug exposure as determinants of heterogeneity in single-cell drug responses. Recent studies have shown that non-cancer cells exhibit variable responses to identical stimuli due to differences in cell state (such as organelle content, size, or confluency), which are stored within cellular signaling networks to guide decision-making (*98*). Similarly, even within clonal cancer cell populations, pre-drug molecular differences, such as NGFR levels, can predetermine divergent resistance trajectories (*14*, *99*). These findings underscore the role of intrinsic cellular diversity, shaped by prior signaling events or molecular context, in contributing to the heterogeneous behaviors observed in cells during drug treatment. Further investigation into how intrinsically different cell states respond to external drug pressures is essential for better understanding cell-fate decisions and identifying strategies to combat drug resistance.

In summary, our work elucidates how persister cells non-genetically adapt to escape drug-induced quiescence and resume proliferation. Notably, we find that cells exhibit varying proliferative potentials under drug treatment, which relate to their ability to reactivate ERK and mTORC1 signaling and ultimately accumulate sufficient Cyclin D1 levels for cell-cycle re-entry. Cyclin D1 therefore represents a potential therapeutic target, where its inhibition, in combination with MAPK inhibitors, could more effectively prevent tumor relapse.

## MATERIALS & METHODS

### Cell culture

The A375 BRAF^V600E^ malignant human melanoma cell line (RRID: CVCL_0132) was purchased from American Type Culture Collection (ATCC). A375 cells were cultured at 37 °C with 5% CO2 in DMEM (Thermo Fisher, #12800-082) supplemented with 10% FBS, 1.5-g/L sodium bicarbonate (Fisher Chemical, #S233-500) and 1X penicillin/streptomycin. The WM278 BRAF^V600E^ malignant melanoma cell line (RRID: CVCL_6473) was obtained from Dr. Natalie Ahn (University of Colorado Boulder). The SKMEL28 BRAF^V600E^ malignant melanoma cell line (RRID: CVCL_0526) was obtained from Dr. Neal Rosen (Memorial Sloan Kettering Cancer Center). WM278 and SKMEL28 cells were maintained at 37°C with 5% CO_2_ in RPMI1640 (Thermo Fisher, #22400-089) supplemented with 10% FBS, 1X Glutamax, 1X sodium pyruvate (Thermo Fisher, #11360-070) and 1X penicillin plus streptomycin. The A375 and WM278 lines were authenticated by short tandem repeat profiling.

### Stable cell line generation

Low-passage A375 cells were transduced with H2B-mIFP lentivirus for cell tracking and DHB-mCherry lentivirus to monitor CDK2 activity, as previously described (*41*). 2-color A375 cells were transduced with ERK KTR-mClover lentivirus to measure ERK activity, prepared with the pLenti-ERKTR-mClover plasmid (*40*). Transduced cells were sorted by fluorescence activated cell sorting (FACS) on the fluorescent colors they carry to establish the cell line.

Integration of the mCitrine-encoding gene into the CCND1 locus of A375 cells was carried out using CRISPR technology. A CRISPR-Cas9 ribonucleoprotein (RNP) complex was generated using the CRISPR-Cas9 System from IDT. The RNP contains crRNA (GGAGCUGGUGUUCCAUGGCUGUUUUAGAGCUAUGCU) annealed to tracrRNA and Cas9 nuclease. The RNP was electroporated into A375 cells using the Neon transfection system (Invitrogen, #MPK10096) with 2 pulses x 20 ms pulse width at 1200 V. Single cells were sorted by FACS into 96-well plates and grown into clonal populations. Western blot and IF against Cyclin D1 protein, PCR of the *CCND1* gene, and genomic sequencing were carried out as validation. Data from clone F4 is shown in this work (Fig. S4 D-F; Data file S1). For functional validation, cells were treated with the MEK inhibitor trametinib, (MedChemExpress, #HY-10999) at 10 nM for 24 hours to observe the decrease in tagged Cyclin D1, and with MLN4924 (Active Biochem, A-1139) at 1.4 μM for 6 hours to block Cyclin D1 degradation and observe its stabilization (Fig. S4E). Validated clone F4 was further multiplexed with H2B-mIFP lentivirus for cell tracking and DHB-mCherry lentivirus as a CDK2 activity sensor in live-cell imaging experiments.

### Small molecules/inhibitors

Treatments used in this study include: dabrafenib (BRAFi, MedChemExpress, #HY-14660); trametinib (MEKi, MedChemExpress, #HY-10999); SCH772984 (ERKi, SelleckChem #S7101); PLX8394 (BRAF paradox breaker, MedChemExpress, #HY-18972); Tovorafenib (Pan-Raf, MedChemExpress, #HY-15246); RapaLink-1 (mTORC1i, MedChemExpress, #HY-111373); Palbociclib (CDK4/6i, Selleckchem #S1116), MLN4924 (NEDD8i, Active Biochem, #A-1139); Chloroquine (autophagy inhibitor, Sigma-Aldrich, #AAJ6445914); 4EGI-1 (translation initiation inhibitor, MedChemExpress, #HY-19831); GDC-0942 (Pictilisib) (PI3Ki, MedChemExpress, #HY-50094). Doses of inhibitors used throughout the study, unless otherwise specified, are: dabrafenib, 1 μM; trametinib, 10 nM; SCH772984, 1 μM; PLX83954, 5 μM; Tovorafenib, 10 μM; RapaLink-1, 100nM; Palbociclib, 1 μM; MLN4924, 1.4 μM; Chloroquine, 10 μM, 4EGI-1, 25 μM, GDC-0942, 3 μM.

### siRNA transfection

siRNA transfections were performed on A375 cells using the DharmaFECT 4 reagent (Dharmacon, #T-2004-02) according to the manufacturer’s instructions. The transfection mix +/-dabrafenib was added to cells after 48 h in dabrafenib or DMSO and removed 6 h later. Cells were then cultured in dabrafenib or DMSO for another 18 hours. Knockdown efficiency was determined by immunofluorescence staining for Cyclin D1 at the time of fixation 72 h after drug and 24 h after transfection. Oligonucleotides were synthesized by Dharmacon: CCND1 (MU-003210-05-0002) or by Horizon Discovery: Negative Control siRNA (D-001810–02-05).

### Antibodies

Primary antibodies and dilutions used for this study include: phospho-Rb (Ser807/811) (D20B12) rabbit mAb (1:500 IF, CST #8516); phospho-Rb (Ser780) mouse mAb (1:1000 IF, BD Biosciences #558385); phospho-ERK1/2 (Thr202/Tyr204) rabbit mAb (1:500 IF, CST #4370); phospho-ERK1/2 (Thr202/Tyr204) mouse mAb (1:1000 IF, MilliporeSigma #M9692); phospho-S6 (Ser240/244) rabbit mAb (1:500 IF, CST #2215); phospho-eIF4e (Ser209) (EP2151Y) rabbit mAb (1:400 IF, Abcam, #ab76256); phospho-4EBP1 (Thr37/46) rabbit mAb (1:250 IF CST # 9456S); LAMP1 D2D11 XP rabbit mAb (1:1000 IF, CST #9091); LC3 II rabbit mAb (1:1000 IF, Abcam, #ab192890); Anti-Cyclin D1 clone SP4 rabbit mAb (1:500 IF Thermo Fisher, RM-9140-S0); Histone H3 mouse mAb (1:2000 WB, CST, #3638); phospho-AKT (Ser473) rabbit mAb (1:200 IF, CST, #9271). Secondary antibodies in this study include goat anti-rabbit, or goat anti-mouseIgG secondaries linked to Alexa Fluor 488 (1:1000 IF, Thermo Fisher #A-11034, #A-11001), Alexa Fluor 546 (1:1000 IF, Thermo Fisher #A-11010, #A-11003, #A-21098), Alexa Fluor 647 (1:1000 IF, Thermo Fisher #A-21245, #A-21235), or HRP (1:4000 for western blots, Cell Signaling Technology #7074, #7076). Unless otherwise noted, all primary and secondary antibody solutions were prepared in 3% BSA blocking buffer at the indicated dilutions.

### Western blotting

Cells were incubated and lysed on ice for 10 min in RIPA lysis buffer supplemented with reducing agent and 1X phosphatase and protease inhibitor (MilliporeSigma #4906845001, #5892970001). Collected lysates were resuspended by vortex and subjected to sonication for 2 min followed by centrifugation at 20,000rcf at 4°C for 20 min. Protein content was determined via Pierce BCA Protein Assay Kit (#23227). Lysates were prepared for SDS-PAGE analysis at 25µg per sample using NuPAGE LDS Sample Buffer (#NP0007) and heated at 95 °C for 5 min. Proteins were separated by Bolt 4-12% Bis-Tris Plus gel (Thermo Fisher, NW04125) and transferred to iBlot 2 PVDF Regular Stacks (Thermo Fisher Scientific, #IB24001) using the iBlot 2 transfer system. The membrane was incubated in 3% BSA (GoldBio, #A-421-250) supplemented with 0.1% Tween-20 (Thermo Fisher, #9005-64-5) at room temperature for 2 h. Primary antibodies were then added and incubated overnight at 4°C in 0.1X PBS/Casein with 0.1% Tween-20 (Fisher, #BP-337100). Membranes were washed with PBS + 0.1% Tween-20 three times for 5 min and then incubated with anti-rabbit or anti-mouse IgG, HRP-linked secondary antibodies for 1 h. The chemiluminescent signals were detected on an Azure C600 and images were analyzed in Fiji.

### Immunofluorescence imaging

Cells were seeded on a glass-bottom 96-well plate (Cellvis, #P96-1.5H-N) coated with collagen (1:50, Advanced Biomatrix #5005) 24 h prior to drug treatment. Seeding densities for 72 h fixed-cell experiments were optimized to prevent contact inhibition by multi-day time points: A375, 1,000 cells/well; WM278, 1,500 cells/well; SKMEL28, 2,000 cells/well. Following treatments, cells were fixed with 4% paraformaldehyde for 10 min at room temperature. Cells were then permeabilized with either 0.1% Triton X-100 at 4°C for 15 min or 100% methanol at room temperature for 10 min, based on optimized antibody staining. Cells were PBS washed 3X and blocked in 3% BSA solution for 1 h at room temperature before incubating with primary antibodies overnight at 4°C. Cells were then washed with PBS three times, and secondary antibodies were added for 2 h at room temperature. Nuclei/DNA content was labelled with Hoechst 33342 (1:10,000) for 5 min and cytoplasm/protein content was labelled with succinimidyl ester 488A (Sigma-Aldrich, #SCJ4600018) (1:10,000 in PBS) for 30 min. 2D-imaging of markers was performed on a Nikon Ti-E using a 10X/0.45 numerical aperture (NA) objective. 3D-imaging of cell cultures for nuclear and cytoplasmic volume measurements was performed on a Nikon AXR laser scanning confocal microscope equipped with a 60X water immersion 1.2 NA objective, Piezo Z-drive, and tunable GaAsp photomultiplier tube detectors. For each cell, a 11.5 µm z-stack was acquired with 0.1 µm spacing using bidirectional resonance scanning.

### Immunofluorescence Image Processing

Image processing for all 2D and 3D images was performed using Fiji and MATLAB Mathworks R2022b, as previously described (*100*). 3D images were pre-processed using Lucy-Richardson confocal deconvolution. Single-cell quantification of captured images was conducted using custom MATLAB scripts. Nuclear signals (p-Rb, p-eiF4E, p-ERK, etc) were quantified from a nuclear mask (median nuclear intensity), which was generated using Otsu’s method on cells stained for Hoechst 33342 signal. Cytoplasmic signals (LAMP1, p-S6, p-ERK, etc) were quantified from a cytoplasmic mask (median cytoplasmic intensity), which was generated using Otsu’s method on cells stained for succinimidyl ester 488A. To measure signal intensity of these markers in escapees vs non-escapees, p-Rb was separated into high and low modes by using the saddle-point in the data as the cutoff. The region props function in MATLAB was used to quantify the median signal for each stain from these masks.

### Measurement of mRNA by fluorescence in situ hybridization (FISH)

Cells were seeded on a glass-bottom 96-well plate coated with collagen 24 h prior to drug treatment. Cells were fixed with 4% paraformaldehyde, and when applicable, were processed for RNA-FISH analysis according to the manufacturer’s protocol (ViewRNA ISH Cell Assay Kit) (Thermo Fisher, #QVC0001). mRNA probes for CCND1 RNA FISH (Thermo Fisher, Type 6 VA6–16943) were hybridized at 40°C for 4 h, followed by standard amplification and fluorescent labelling steps also at 40 °C. Blocking and immunofluorescence staining was performed after FISH labeling. FISH images were taken on PerkinElmer Opera Phenix high-content screening system with a 20x/1.0 NA water objective, and puncta detection was performed using Harmony image analysis software.

### Live-cell time-lapse imaging and single-cell tracking

Cells were seeded on a glass-bottom 96-well plate coated with collagen 24 h prior to the start of imaging. Movie images for each fluorescent channel were taken on a Nikon Ti-E using a 10X/0.45 NA objective (for imaging A375 cells with exogenous sensors) or 20X/0.75 NA objective (for imaging the endogenously tagged A375 mCitrine-Cyclin D1 clone) at a 15 min frame rate. Cells were maintained in a humidified incubation chamber at 37°C with 5% CO2. Cells were imaged in phenol red-free full-growth media. For times of drug addition or siRNA treatment, the movie was paused to allow for exchange of treatment-containing media, and then imaging continued. For treatments spanning multiple days, drug was refreshed 48 h after the initial drug addition time point.

Multi-day movies were tracked and processed using established pipelines, as previously described (*21*, *30*). The open access tracking codes are available for download from GitHub at https://github.com/tianchengzhe/EllipTrack. CDK2 activity was read out as the cytoplasmic: nuclear (C/N) ratio of the DHB signal, as previously described (*100*), where nuclear mean was used and the cytoplasmic component was calculated as the mean of the top 50th percentile of a ring of pixels outside of the nuclear mask. The cell-cycle entry point (Restriction point) is defined as the point where CDK2 activity begins to rise. The point of anaphase is defined when the H2B-based nuclear mask is divided to give rise to two new distinctly segmented bodies. Escapees are defined as cells with at least 10 h of drug-induced CDK2^low^ quiescence before cell-cycle re-entry. Non-escapees are defined as cells that never emerge from drug-induced CDK2-low quiescence before the end of the imaging period. For CDK2 activity heatmaps, escapee and non-escapee lineages were plotted separately and cells within each category were sorted by genealogy emergence. The plots were then combined. Lineages included for plotting had at least one mitosis prior to drug-induced quiescence and were tracked for the complete duration of drug treatment. Vistrack was used to verify complete and accurate tracking of at least three sites throughout the imaging period of every movie acquired and analyzed.

### Statistics

For violin plots used throughout the paper, thick lines represent the median values unless otherwise indicated. The full distributions are displayed by the full range of the violin shape, with the width along the violin corresponding with the value frequency. Data plotted throughout are representative of at least 3 independent experiments for each figure panel. Error bars in our figures indicate the standard error of the mean, which was calculated from multiple technical replicates. Statistical tests were performed using GraphPad Prism and MATLAB, and test type was chosen based on sample size: n = 3-10, unpaired t-test (parametric); n > 3-10, Mann-Whitney U-test (nonparametric). Significance levels are reported as *p* values ≤ 0.05 (*), 0.01 (**), 0.001 (***) and 0.0001 (****) with corresponding star notations. Throughout, ‘ns’ denotes no statistical significance (*p* > 0.05). All instances of log refer to the natural log, which is the default in MATLAB.

### Receiver operating characteristic (ROC) analysis

For all growth rate ROC analyses, we determined the false positive (non-escapees) and true positive (escapees) rate of detection for the total growth rate during the CDK2^low^ period prior to cell-cycle re-entry by sliding the cutoff at every 10^th^ percentile of rate. Our classification of true-positive escapee cells required that cells had at least 10 h of drug-induced CDK2^low^ quiescence before cell-cycle re-entry. For all Cyclin D1 ROC analyses, escapees that were CDK2^low^ for at least 15 h before escaping were aligned to their R point and the Cyclin D1 intensity at different timepoints (–5, –10, and –15 h) prior to cell-cycle re-entry was obtained for threshold detection. For every escapee cell, a non-escapee cell was randomly chosen and aligned to the R point timing of an escapee cell, enabling acquisition of Cyclin D1 intensity. The false positive (non-escapees) and true positive (escapees) rate of detection was again determined for Cyclin D1 intensity by sliding the cutoff at every 10^th^ percentile of intensity.

## Supplementary Materials

**Fig. S1.** Cycling persister cells are observed across multiple BRAF^V600E^-mutant melanoma lines, related to Fig. 1.

**Fig. S2.** ERK activity is highly dynamic and heterogenous over time in drug-treated A375 cells related to Fig. 2.

**Fig. S3.** mTORC1-mediated growth and mass accumulation influences cell-cycle re-entry across multiple BRAF^V600E^-mutant melanoma lines, related to Fig.4.

**Fig. S4.** Manipulations to Cyclin D1 protein levels and characterization of the mCitrine-Cyclin D1 knock-in A375 cell line, related to Fig. 5.

**Fig. S5.** Using the mCitrine-Cyclin D1 A375 line to study Cyclin D1 dynamics in the presence of MAPK pathway inhibitors, related to Fig. 5.

**Fig. S6.** PI3K/AKT signaling is not the dominant mode of mTORC1 activation in melanoma cell lines escaping dabrafenib.

**The Data file S1**. Genome sequencing of the Cyclin D1 locus in A375 cells after mCitrine-Cyclin D1 knock-in.

**Movie S1**. Correlating CDK2 activity sensor and mCitrine-Cyclin D1 levels in dabrafenib-treated A375 cells

## Acknowledgements

We thank members of the Spencer Lab for general help and discussion. We thank Theresa Nahreini, head of the Biochemistry Cell Culture Facility (BCCF), for her expertise and assistance with cell sorting. The BD FACSCelesta cytometer and BD FACSAria Fusion cell sorter are supported by NIH Grant S10ODO21601. We thank Joe Dragavon, head of the BioFrontiers Advanced Light Microscopy Core, for his insights on quantitative microscopy and aid in image acquisition. The PerkinElmer Opera Phenix is supported by NIH grant 1S10OD025072. 3D-confocal microscopy was performed on a Nikon AXR laser scanning confocal microscope supported by the NIST-CU Cooperative Agreement award number 70NANB15H226.

## Author contributions

V.N. designed and conducted experiments, analyses, data interpretation and manuscript preparation. N.M. created the CCND1-mCitrine A375 cell line and N.M and V.N. performed validation experiments. V.N., H.A., and V.P. developed analysis scripts for data analysis. V.P. imaged and analyzed cell volume measurements using confocal imaging. H.A. and C.R.I. contributed to experimental design. S.L.S. conceived the project with intellectual contributions from V.N. throughout. V.N and S.L.S wrote the manuscript. S.L.S. acquired funding and supervised the project.

## Data and materials availability

Raw source data and materials used in this study are available upon request.

## Code availability

All custom MATLAB scripts are available upon request. Single-cell tracking scripts and analysis scripts are available in GitHub at: https://github.com/tianchengzhe/EllipTrack. Immunofluorescence imaging processing and analysis scripts are available in GitHub at: https://github.com/tiho9814/matlab_image_processing_scripts/tree/main/scripts_IF.

