## Supplement for "MAPK and mTORC1 signaling converge on Cyclin D to enable cell-cycle re-entry in melanoma persister cells"

**<u>Supplementary Materials</u>**

Figures S1 to S6

Data file S1

Movie S1

### Supplementary Figures:

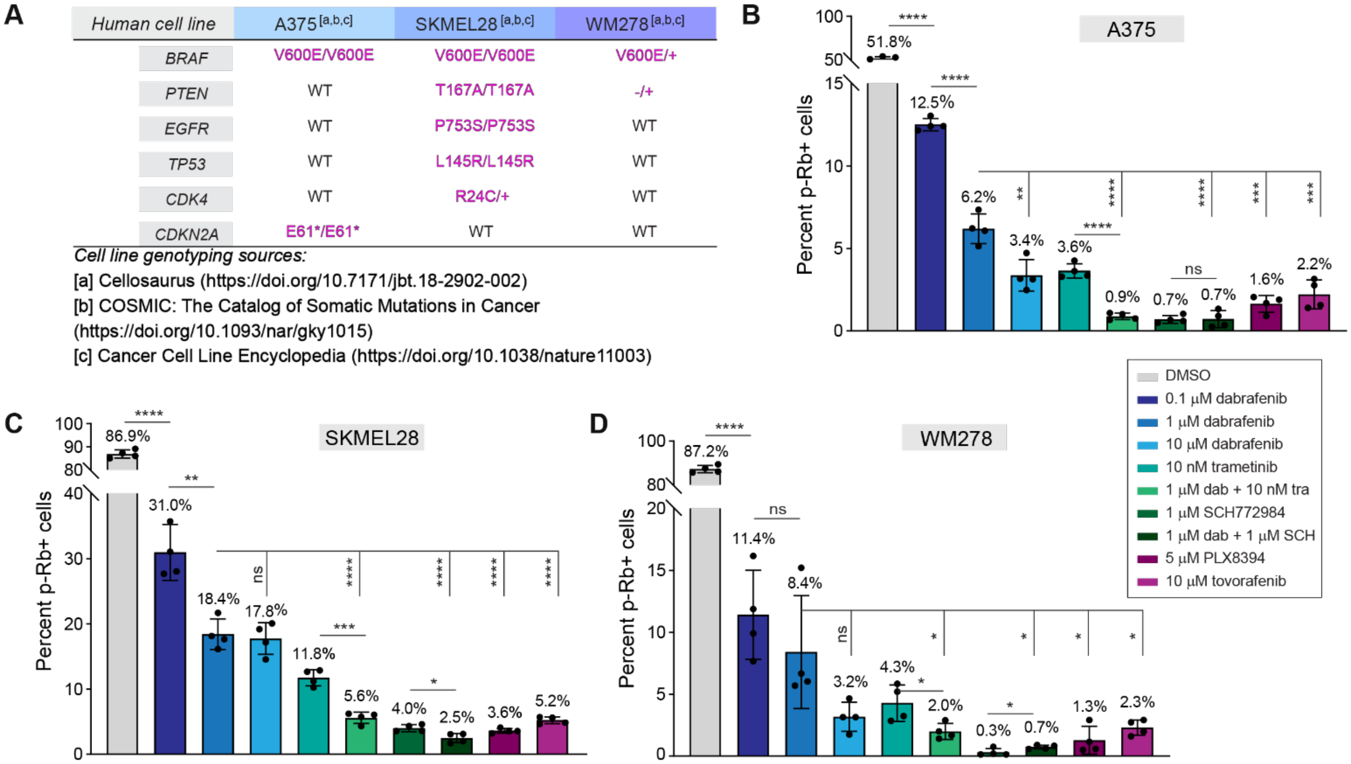


**Figure. S1 | Cycling persister cells are observed across multiple BRAF$^{V600E}$-mutant melanoma lines, related to** Fig. 1**.**

**(A)** Genotype of each human BRAF$^{V600E}$-mutant melanoma cell line identified as reported by indicated databases. **(B-D)** Profiling of percentage of cycling cells (p-Rb$^{+}$) 72 h after treatment with the indicated MAPK pathway inhibitor in A375 (B), SKMEL28 (C), and WM278 cells (D). Data are mean ± std of 4 replicate wells imaged, representative of 3 independent experiments. For bar graphs, *p*-values were calculated by unpaired t-tests between indicated drug conditions, specified with a black line linking the conditions.

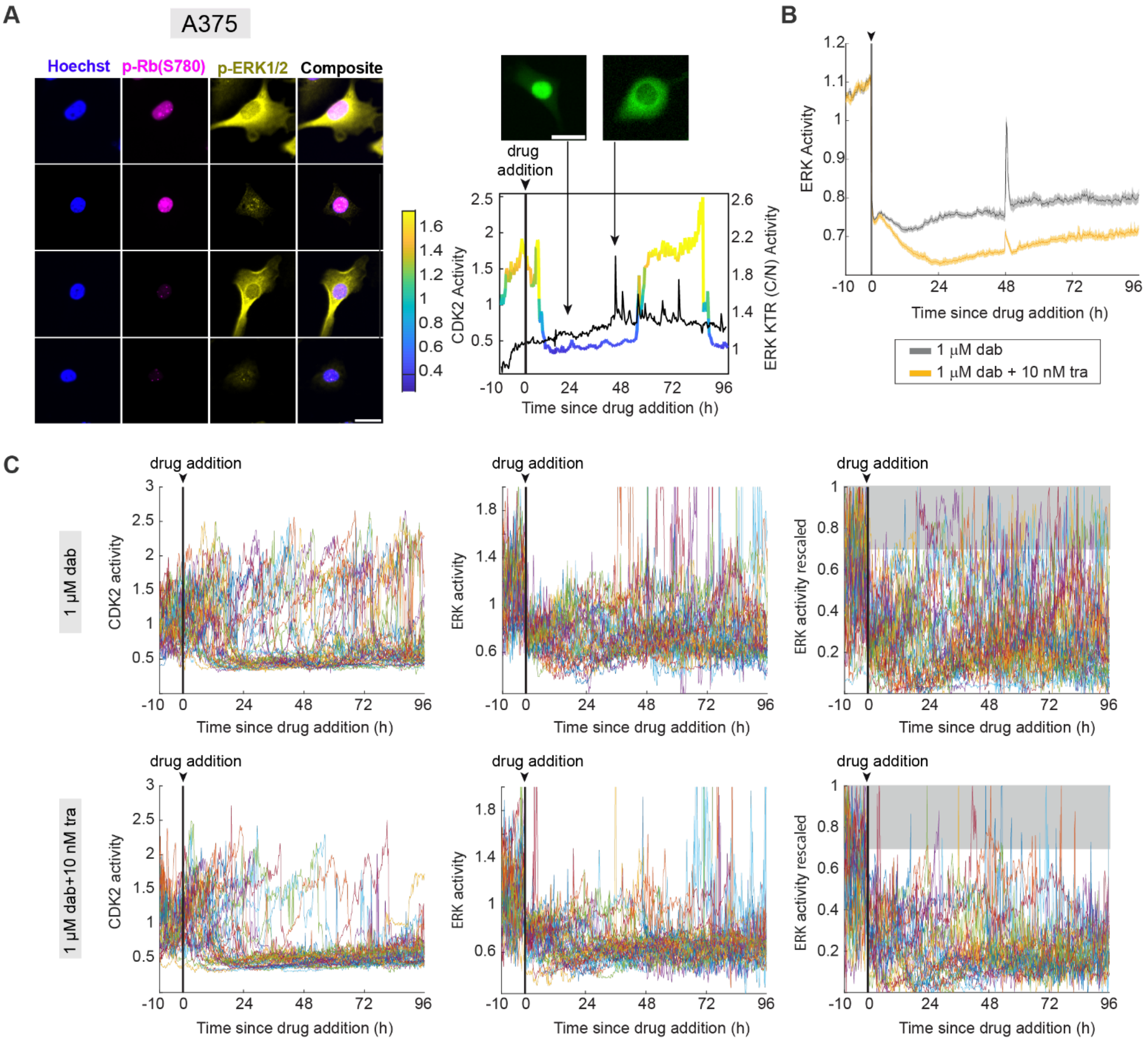


**Figure. S2 | ERK activity is highly dynamic and heterogenous over time in drug-treated A375 cells related to** Fig. 2**.**

**(A) Left:** Representative IF images of A375 cells co-stained for Hoescht, p-Rb (S780), and p-ERK1/2 (T202/Y204) after 1 µM dabrafenib for 72 h. Note that at a single snapshot in time, both cycling (p-Rb$^{+}$) and non-cycling cells (p-Rb$^{-}$) can have high ERK activity or low ERK activity. Scalebar = 30 µm. **Right:** Representative single-cell trace of CDK2 activity (parula colormap) and ERK KTR$_{(C/N)}$ activity (black) in a single A375 cell treated with 1 µM dabrafenib to highlight ERK's dynamic and variable temporal activity. Trace is overlaid with representative images from the ERK KTR sensor demonstrating how fixing and staining for p-ERK1/2 is limited due to ERK's heterogenous temporal dynamics. Scalebar = 25 µm**.** **(B)** Population average and 95% confidence interval of temporal ERK activity from ERK KTR sensor in A375 cells treated for 72 h with either 1 µM dabrafenib (gray) or combination 1 µM dabrafenib + 10 nM trametinib (yellow). The spike at 48 h is the result of a media refresh. **(C)** 40 randomly selected A375 single cell traces showing temporal CDK2 activity measured by CK2 activity sensor (DHB-mCherry) (left), temporal ERK activity from ERK KTR sensor *(*middle), and the same temporal ERK activity rescaled across the minimum and maximum value for each individual cell in order to highlight ERK activity pulses.

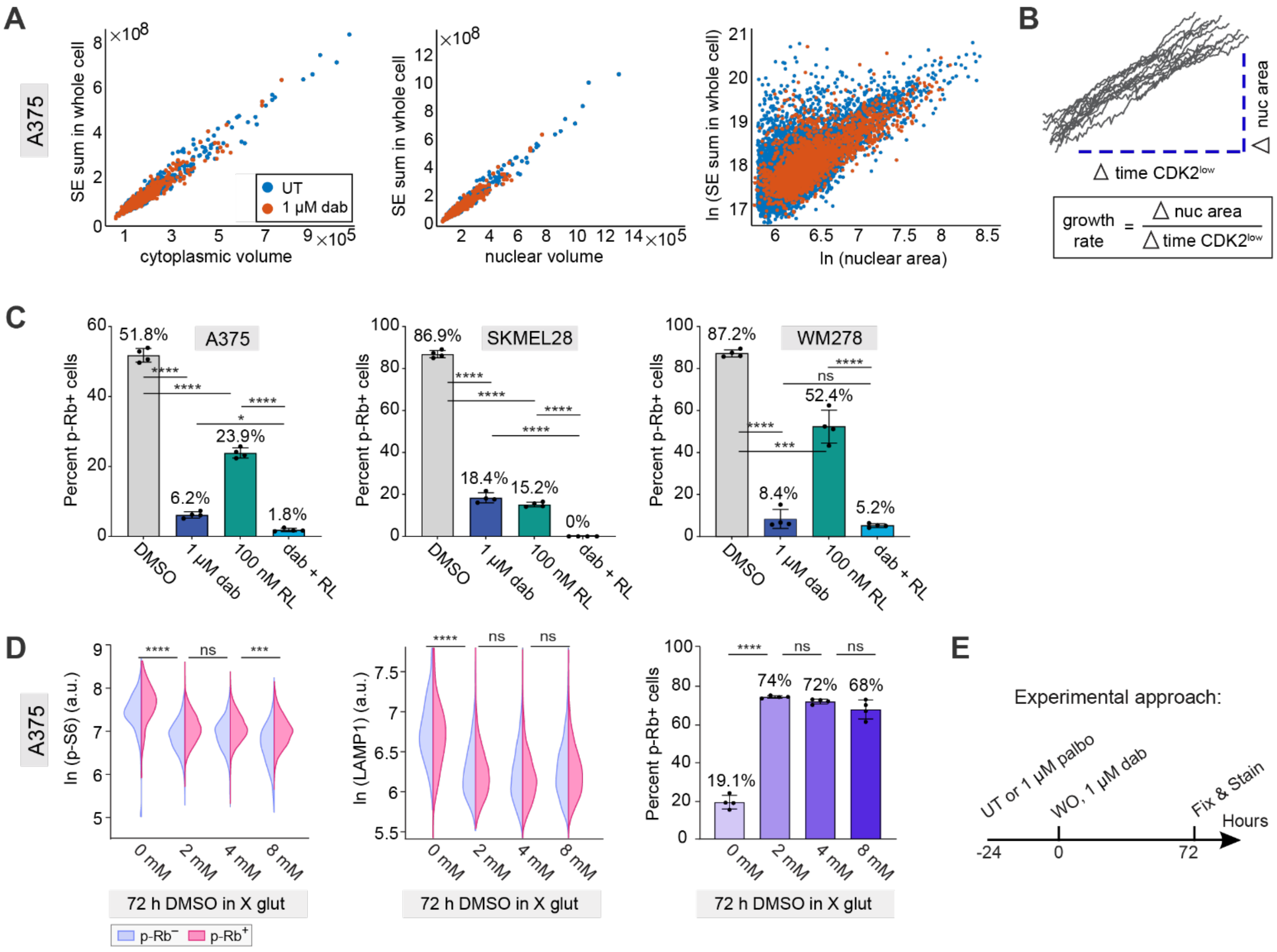


**Figure. S3 | mTORC1-mediated growth and mass accumulation influences cell-cycle re-entry across multiple BRAF$^{V600E}$-mutant melanoma lines, related to Fig.4.**

**(A)** Left: Scatter plots of cytoplasmic volume versus total protein stain succinimidyl ester in each whole cell (nuclear + cytoplasmic volume) obtained from 3D confocal imaging; Middle: Scatter plots of nuclear volume versus succinimidyl ester in each whole cell obtained via 3D confocal imaging; Right: Scatter plots of nuclear area versus succinimidyl ester in whole cell (nuclear + cytoplasmic area) obtained via 2D epiflouresence microscopy. **(B)** Schematic representing growth rate quantification as the change in nuclear area only during the quiescent CDK2$^{low}$ period (>0.8 cutoff) prior to cell-cycle re-entry. **(C)** Profiling of the percentage of cycling cells (p-Rb$^{+}$) 72 h after treatment with either DMSO, 1 µM dabrafenib, 100 nM RapaLink-1, or the combination in A375, SKMEL28, and WM278 cells. **(D)** A375 cells treated with 1 µM dabrafenib for 72 h in increasing concentrations of glutamine (standard concentration is 4 mM) and stained for p-S6 (S240/244) (left), LAMP1 (middle), and p-Rb (S780) (right). For split violins, *p*-values were determined by Mann-Whitney U-tests between escapees (p-Rb$^{+}$) from the indicated drug conditions, specified with a black line. **(H)** Schematic of experimental drug conditions with associated legend. **(E)** Schematic of experimental approach for Fig. 4, G and H. WO, washout. For bar graphs, *p*-values were calculated by unpaired t-tests between indicated drug conditions, specified with a black line linking the conditions. Error bars: mean ± std of four replicate wells.

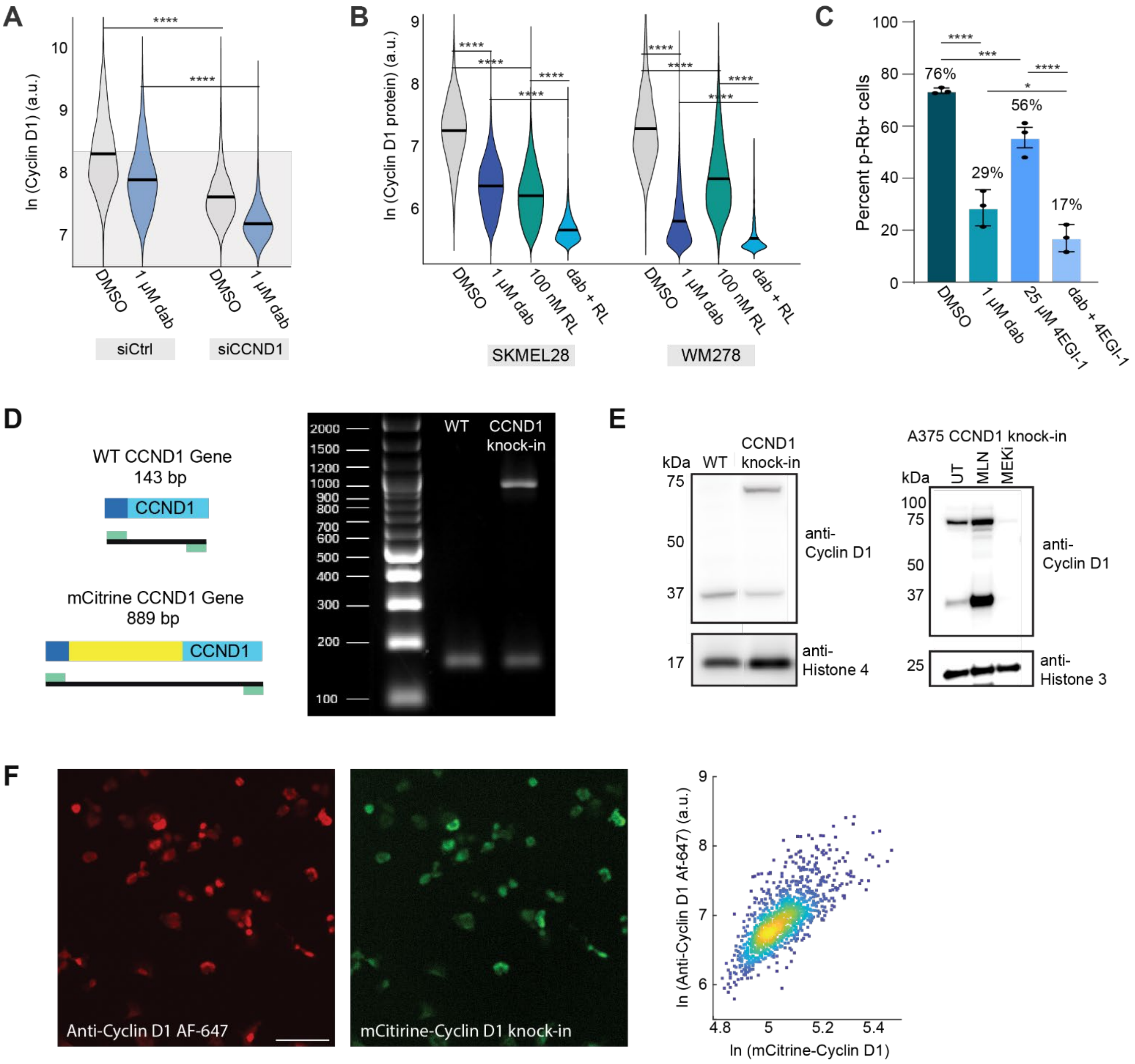


**Figure. S4 | Manipulations to Cyclin D1 protein levels and characterization of the mCitrine-Cyclin D1 knock-in A375 cell line, related to Fig. 5.**

**(A)** Bulk violin plots quantifying the reduction in Cyclin D1 protein, obtained from IF staining, after 72 h of treatment with either DMSO or 1 μM dabrafenib and 24 h after transfection with siControl or siCCND1 knockdown constructs. Transfection mixture +/- dabrafenib was added to cells after 48 h in dabrafenib or DMSO and removed 6 h later. Cells were then cultured in dabrafenib or DMSO for another 18 hours. **(B)** Bulk violins of Cyclin D1 protein, quantified by IF, in A375 cells treated for 72 h with either DMSO, 1 μM dabrafenib, 100 nM RapaLink-1, or the combination in SKMEL28, and WM278 cells. Note that Cyclin D1 protein levels directly correlate to the percentage of cycling cells under the same conditions in Fig. S3C. **(C)** Percentage of cycling A375 cells (p-Rb$^{+}$) 72 h after treatment with either DMSO, 1 μM dabrafenib, 25 μM 4EGI-1 (translation initiation inhibitor), or the combination. **(D)** Left: The expected DNA band sizes for the wild-type CCND1 gene and the mCitrine-Cyclin D1 fusion. Right: PCR amplification of the CCND1 gene in parental and mCitrine-CCND1 knock-in cells run on 0.8% agarose gel. The mCitrine gene was heterozygously knocked into only one allele resulting in the presence of a wild-type band in the mCitrine-CCND1 knock-in cells. Bands were excised and sequenced as additional verification (see also Data file S1). **(E)** Left: Western blot of parental and mCitrine-CCND1 knock-in A375 cells run for molecular weight verification. Right: The mCitrine-Cyclin D1 knock-in A375 cell responds in the same way as untagged Cyclin D1 in parental A375 cells (increased by MLN4924 and decreased by Meki). WT: parental A375 cells. CCND1 knock-in: A375 cells with mCitrine knocked into the CCND1 locus to produce a mCitrine-Cyclin D1 fusion protein. UT: Untreated. MLN: 1.4 µM MLN4924 treatment for 6 hours. Meki: 100 nM PD0325901 treatment for 24 h. **(F)** Left: representative IF images of mCitrine signal and Cyclin D1 antibody staining in the same cells. Right: Quantification of images showing mCitrine intensity linearly correlates with Cyclin D1 antibody staining in the mCitrine-Cyclin D1 knock-in cell line. Scalebar =100 µm. For bar graphs, *p*-values were calculated by unpaired t-tests between indicated drug conditions, specified with a black line linking the conditions. Error bars: mean ± std of four replicate wells. For bulk violins, *p*-values were determined by Mann-Whitney U-tests between indicated drug conditions, specified with a black line linking the conditions.

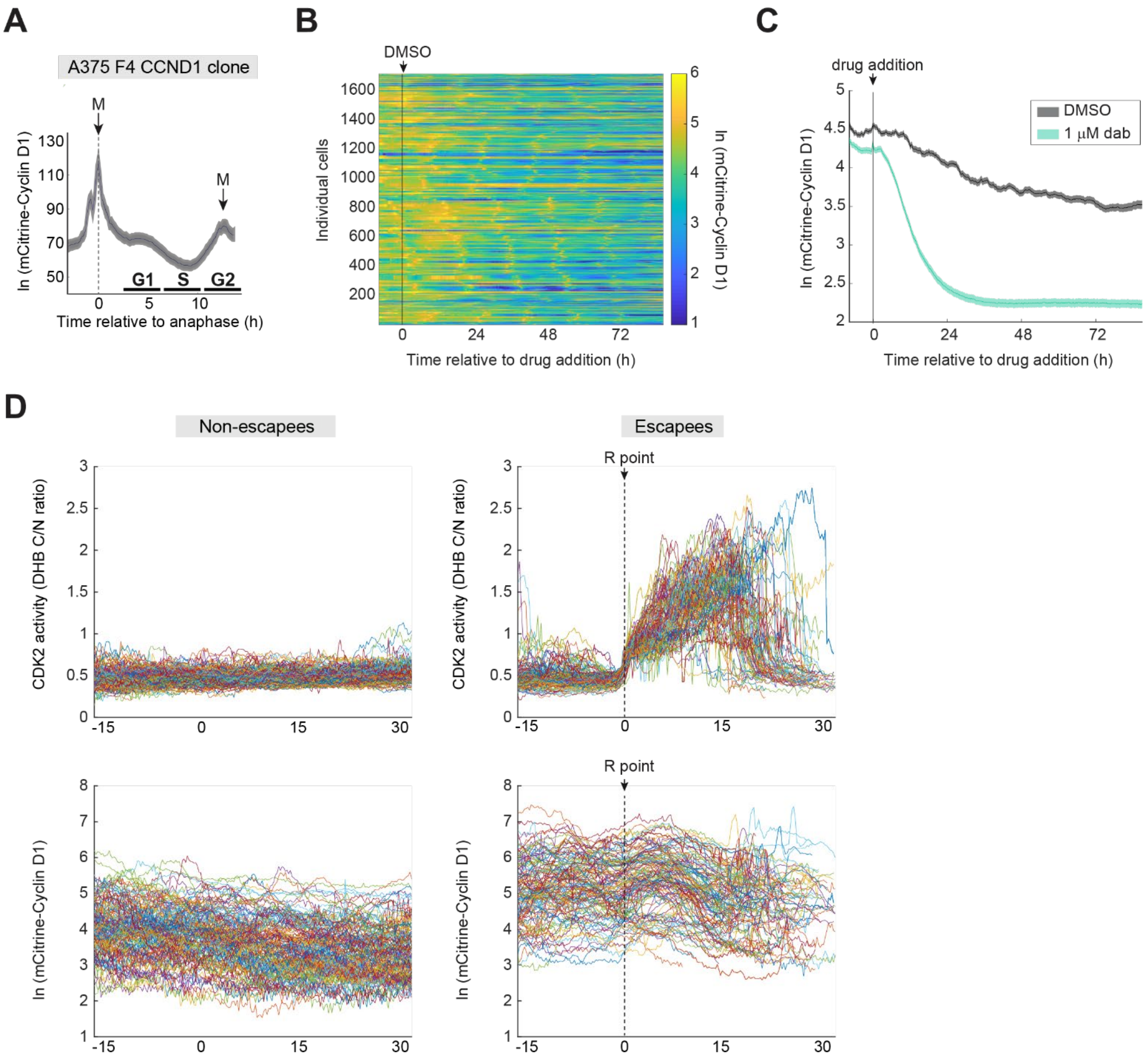


**Figure. S5 | Using the mCitrine-Cyclin D1 A375 line to study Cyclin D1 dynamics in the presence of MAPK pathway inhibitors, related to Fig. 5.**

**(A)** Population average and 95% confidence interval of temporal CCND1 levels in the A375 mCitrine-Cyclin D1 knock-in cells treated with either DMSO or 1 μM dabrafenib. Note that upon S phase entry, Cyclin D1 gets degraded, as previously reported (*25*, *78*). **(B)** Heatmap of temporal endogenous mCitrine-Cyclin D1 in thousands of single A375 cells after DMSO addition at the arrow and black line mark. Each row represents the endogenous Cyclin D1 levels in a single cell over time according to the colormap. Sample size is defined on y-axis. **(C)** Population average and 95% confidence interval of mCitrine-Cyclin D1 levels in the A375 mCitrine-Cyclin D1 knock-in cells treated with either DMSO or 1 μM dabrafenib. Note the rapid reduction in Cyclin D1 protein levels within a few hours of drug addition. **(D)** Single cell traces of CDK2 activity (top) and endogenous mCitrine-Cyclin D1 (bottom) in 1 μM dabrafenib-treated A375 cells for non-escapees (left) or escapees (right) aligned to the restriction point (R point, the point of CDK2 activity rise) in order to visualize endogenous Cyclin D1 levels prior to escape.

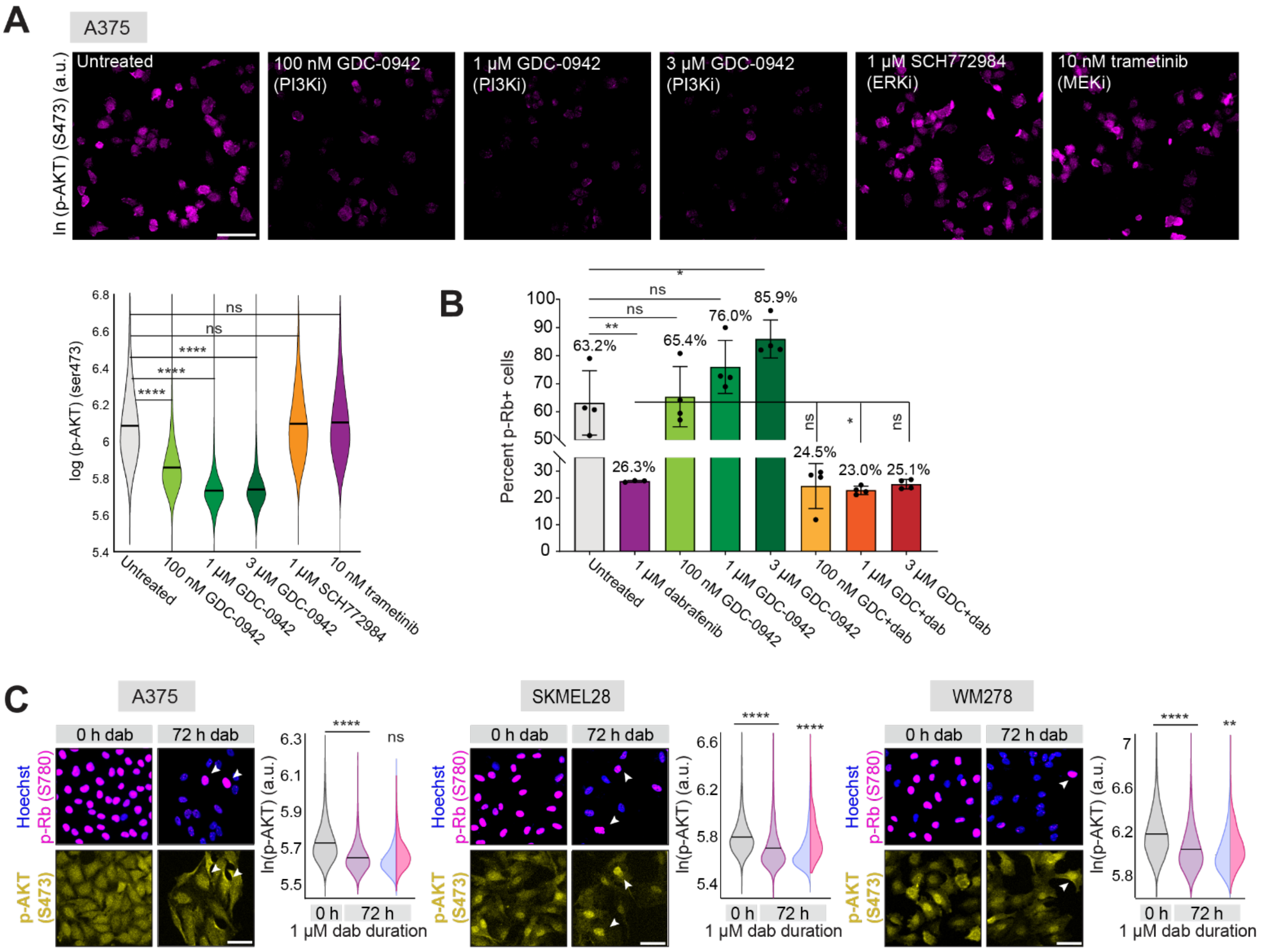


**Figure. S6 | PI3K/AKT signaling is not the dominant mode of mTORC1 activation in melanoma cell lines escaping dabrafenib.**

**(A)** Top: representative IF images of A375 cells stained for AKT activity marker p-AKT (S473) 2 h after treatment with DMSO, increasing concentrations of either the PI3K inhibitor GDC-0941, or the ERK inhibitor SCH772984. Scalebar = 100 µm Bottom: Bulk violin plots quantifying p-AKT levels from representative images above and validating GDC-0941 as a strong and selective PI3K inhibitor. **(B)** Percentage of cycling A375 cells (p-Rb$^{+}$) 72 h after treatment with the indicated drug conditions, showing minimal difference between dabrafenib treatment alone or dabrafenib plus the PI3Ki inhibitor GDC-0941. **(C)** Representative IF images of either A375, SKMEL28, or WM278 cells treated with 1 µM dabrafenib for 72 h and co-stained for Hoescht, p-Rb (S780), and p-AKT (S473). Quantification of each marker is plotted as bulk violin plots in untreated cells (gray) vs dabrafenib-treated (purple) cells as well as split violin plots in dabrafenib-treated cells to highlight differences in activity between non-escapees (p-Rb$^{-}$, blue) vs. escapees (p-Rb$^{+}$, pink). Note that a strong difference in p-AKT activity between escapees and non-escapees is only seen in SKMEL28 cells, which harbor PTEN mutations, as described in Fig S1A. Scalebar = 50 µm. For bar graphs, *p*-values were calculated by unpaired t-tests between indicated drug conditions, specified with a black line linking the conditions. Error bars: mean ± std of four replicate wells. For bulk violins, *p*-values were determined by Mann-Whitney U-tests between indicated drug conditions, specified with a black line linking the conditions.

| | |
|---|---|
| Reference Sequence | Highlighted in pale blue |
| A375 F4 mCitrine-CCND1 Aligned Sequence | Not highlighted |
| Start of mCitrine | |
| Linker | |
| Start of Cyclin D1 Coding Region | |
| End of Cyclin D1 Coding Region | |
| Known SNPs in Cyclin D1 | |
| Exons | Underlined in Reference Sequence |
| Introns | Not Underlined in Reference Sequence |

GTCGGCGCAGTAGCAGCGAGCAGCAGAGTCCGCACGCTCCGGCGAGGGGCAGAAGAGCGCGAGGGAGCGC

GTCCGCACGCTCCGGCGAGGGGCAGAAGAGCGCGAGGGAGCGC

GGGGCAGCAGAAGCGAGAGCCGAGCGCGGACCCAGCCAGGACCCACAGCCCTCCCCAGCTGCCCAGGAAG

GGGGCAGCAGAAGCGAGAGCCGAGCGCGGACCCAGCCAGGACCCACAGCCCTCCCCAGCTGCCCAGGAAG

AGCCCCAGCCATGGTGAGCAAGGGCGAGGAGCTGTTCACCGGGGTGGTGCCCATCCTGGTCGAGCTGGAC

AGCCCCAGCCATGGTGAGCAAGGGCGAGGAGCTGTTCACCGGGGTGGTGCCCATCCTGGTCGAGCTGGAC

GGCGACGTAAACGGCCACAAGTTCAGCGTGTCCGGCGAGGGCGAGGGCGATGCCACCTACGGCAAGCTGA

GGCGACGTAAACGGCCACAAGTTCAGCGTGTCCGGCGAGGGCGAGGGCGATGCCACCTACGGCAAGCTGA

CCCTGAAGTTCATCTGCACCACCGGCAAGCTGCCCGTGCCCTGGCCCACCCTCGTGACCACCTTCGGCTA

CCCTGAAGTTCATCTGCACCACCGGCAAGCTGCCCGTGCCCTGGCCCACCCTCGTGACCACCTTCGGCTA

CGGCCTGATGTGCTTCGCCCGCTACCCCGACCACATGAAGCAGCACGACTTCTTCAAGTCCGCCATGCCC

CGGCCTGATGTGCTTCGCCCGCTACCCCGACCACATGAAGCAGCACGACTTCTTCAAGTCCGCCATGCCC

GAAGGCTACGTCCAGGAGCGCACCATCTTCTTCAAGGACGACGGCAACTACAAGACCCGCGCCGAGGTGA

GAAGGCTACGTCCAGGAGCGCACCATCTTCTTCAAGGACGACGGCAACTACAAGACCCGCGCCGAGGTGA

AGTTCGAGGGCGACACCCTGGTGAACCGCATCGAGCTGAAGGGCATCGACTTCAAGGAGGACGGCAACAT

AGTTCGAGGGCGACACCCTGGTGAACCGCATCGAGCTGAAGGGCATCGACTTCAAGGAGGACGGCAACAT

CCTGGGGCACAAGCTGGAGTACAACTACAACAGCCACAACGTCTATATCATGGCCGACAAGCAGAAGAAC

CCTGGGGCACAAGCTGGAGTACAACTACAACAGCCACAACGTCTATATCATGGCCGACAAGCAGAAGAAC

GGCATCAAGGTGAACTTCAAGATCCGCCACAACATCGAGGACGGCAGCGTGCAGCTCGCCGACCACTACC

GGCATCAAGGTGAACTTCAAGATCCGCCACAACATCGAGGACGGCAGCGTGCAGCTCGCCGACCACTACC

AGCAGAACACCCCCATCGGCGACGGCCCCGTGCTGCTGCCCGACAACCACTACCTGAGCTACCAGTCCAA

AGCAGAACACCCCCATCGGCGACGGCCCCGTGCTGCTGCCCGACAACCACTACCTGAGCTACCAGTCCAA

GCTGAGCAAAGACCCCAACGAGAAGCGCGATCACATGGTCCTGCTGGAGTTCGTGACCGCCGCCGGGATC

GCTGAGCAAAGACCCCAACGAGAAGCGCGATCACATGGTCCTGCTGGAGTTCGTGACCGCCGCCGGGATC

ACTCTCGGCATGGACGAGCTGTACAAGTCGGGCGGATCCGGCGAACACCAGCTCCTGTGCTGCGAAGTGG

ACTCTCGGCATGGACGAGCTGTACAAGTCGGGCGGATCCGGCGAACACCAGCTCCTGTGCTGCGAAGTGG

AAACCATCCGCCGCGCGTACCCCGATGCCAACCTCCTCAACGACCGGGTGCTGCGGGCCATGCTGAAGGC

AAACCATCCGCCGCGCGTACCCCGATGCCAACCTCCTCAACGACCGGGTGCTGCGGGCCATGCTGAAGGC

GGAGGAGACCTGCGCGCCCTCGGTGTCCTACTTCAAATGTGTGCAGAAGGAGGTCCTGCCGTCCATGCGG

GGAGGAGACCTGCGCGCCCTCGGTGTCCTACTTCAAATGTGTGCAGAAGGAGGTCCTGCCGTCCATGCGG

AAGATCGTCGCCACCTGGATGCTGGAGGTGCGGGGCTTCGGGCGGCTCTCTTAAGACTTCCCTGCAACTT

AAGATCGTCGCCACCTGGATGCTGGAGGTGCGGGGCTTCGGGCGGCTCTCTTAAGACTTCCCTGCAACTT

GTTGCCCAGACCCACGTTTCTTTGCTACTCACCCCCCTCCCTTCTCTCCCGCTAGAACTTTGAAGTTTGC

GTTGCCCAGACCCACGTTTCTTTGCTACTCACCCCCCTCCCTTCTCTCCCGCTAGAACTTTGAAGTTTGC

CGTGGTGTTTCTAGGGATCCGTATTTTCAAAATAAAAATTGCGGGTATTTTCTGAAGGAGGAAGGGGTGG

CGTGGTGTTTCTAGGGATCCGTATTTTCAAAATAAAAATTGCGGGTATTTTCTGAAGGAGGAAGGGGTGG

GGGTGGGGGTGCTAGAAGTAGCGTTTCGTGGGAGGGGAGAAGGGGGTCCGGGAGGGGTGCCTTCGGGAGA

GGGTGGGGGTGCTAGAAGTAGCGTTTCGTGGGAGGGGAGAAGGGGGTCCGGGAGGGGTGCCTTCGGGAGA

AGCCAGTGCCAGGGGCACCCCAATGGGCCCGAGGGTGCGGGCTGGCAGGCTGGGTGCGCTTTGTGTCCCC

AGCCAGTGCCAGGGGCACCCCAATGGGCCCGAGGGTGCGGGCTGGCAGGCTGGGTGCGCTTTGTGTCCCC

CGCCTGCGCCCCAGCCCGGCTGCGCCTCAGCGGCCGGGAGCCGCCAACTCCGGGGGGAGGGGGCATAGAT

CGCCTGCGCCCCAGCCCGGCTGCGCCTCAGCGGCCGGGAGCCGCCAACTCCGGGGGGAGGGGGCATAGAT

TTGATTTTTAAATTAATATCCATGGACACGTATGCAAGGGCCGCTCGTGCCAGTATTATGCGCCATCTTT

TTGATTTTTAAATTAATATCCATGGACACGTATGCAAGGGCCGCTCGTGCCAGTATTATGCGCCATCTTT

GCTCTTTTATTGCAAAGCAAAAGTGTTTATTAATAATTGGGGGCAGGGTGGGGGCGGGGAGCGGCCGCCG

GCTCTTTTATTGCAAAGCAAAAGTGTTTATTAATAATTGGGGGCAGGGTGGGGGCGGGGAGCGGCCGCCG

GGCGCTGGGGCCGCAGCTAAGGGCCGCGCGGCTGCCGGGAGCCCGCGGGAGGGGCGCAGGGACGCGGCAT

GGCGCTGGGGCCGCAGCTAAGGGCCGCGCGGCTGCCGGGAGCCCGCGGGAGGGGCGCAGGGACGCGGCAT

GGGTAGTTTTGGGGGGACGCCGCTAGGGAAGGGGGGGCCTTTGTTCAAGCAGCGAGTCCCGGGGCGCCCC

GGGTAGTTTTGGGGGGACCCCGCTAGGGAAGGGGGGGCCTTTGTTCAAGCAGCGAGTCCCGGGGCGCCCC

GAACGGGCAGCCTGGGCCGGAGAGCACGGCGAGCTGCAAGGTCGCGTGGCCCCCAAGACGCCAGGGCTTG

GAACGGGCAGCCTGGGCCGGAGAGCACGGCGAGCTGCAAGGTCGCGTGGCCCCCAAGACGCCAGGGCTTG

ATCCCCGTCTGCAGGGATATCGGCTTGGAGGACCTTCTCCGAGCGAGCCGGGGGCCTGGGAGCACATTTT

ATCCCCGTCTGCAGGGATATCGGCTTGGAGGACCTTCTCCGAGCGAGCCGGGGGCCTGGGAGCACATTTT

CAGACCTTCGGTGGGCGCCTGAGGGGCCCGCAAGTATTTTAAAATAATTTTTGAAAGTGCGGCGTGGTGC

CAGACCTTCGGTGGGCGCCTGAGGGGCCCGCAAGTATTTTAAAATAATTTTTGAAAGTGCGGCGTGGTGC

CCTTGCGAGAGGGAAACGCCGCCCGCGCCCAGGGGGAAGGGGGGGCCCCGGAGTTTGAATTCCTGGGGCT

CCTTGCGAGAGGGAAACGCCGCCCGCGCCCAGGGGGAAGGGGGGGCCCCGGAGTTTGAATTCCTGGGGCT

CCCCCCGGAGCCTGTAACGAACTCCCAACCCCCGGCCTGGGTAAAGGGTCGCCCGAGGGTCATTTTCAGG

CCCCCCGGAGCCTGTAACGAACTCCCAACCCCCGGCCTGGGTAAAGGGTCGCCCGAGGGTCATTTTCAGG

GTTTTTTTATGCACTTAGTTATTTTTTTAATATTTTTAAATATTTTTTGAAAAGATGACGTCTGGGGAAA

GTTTTTTTATGCACTTAGTTATTTTTTTAATATTTTTAAATATTTTTTGAAAAGATGACGTCTGGGGAAA

TGCGGCGCGGCGGCCTGGGACGCCACCTTTGTGTCTCGCAGGCGCGGCGCCCAACCCCGCGGCCCGTTCC

TGCGGCGCGGCGGCCTGGGACGCCACCTTTGTGTCTCGCAGGCGCGGCGCCCAACCCCGCGGCCCGTTCC

GCGGCCCCGCACCCCAGTTGGTGTCGACCCCCAGTCAGAGGGACCACGGAGCTCCAGGGCGGGCCAGGGT

GCGGCCCCGCACCCCAGTTGGTGTCGACCCCCAGTCAGAGGGACCACGGAGCTCCAGGGCGGGCCAGGGT

CCCGGGGGCCGGCAGCCCGCGCCGCCGCGCACGCCGCCCAGCTGTGCCCGCTCCCGCCCCCACCGTGCCA

CCCGGGGGCCGGCAGCCCGCGCCGCCGCGCACGCCGCCCAGCTGTGCCCGCTCCCGCCCCCACCGTGCCA

GCCTCGCGGGGACTTTCCCTTTCAGTTTCGGGGAGGGTGGGTACTGGGGACGCGCGGGGGAGGGGGCGCA

GCCTCGCGGGGACTTTCCCTTTCAGTTTCGGGGAGGGTGGGTACTGGGGACGCGCGGGGGAGGGGGCGCA

TCACGGGAAGCTCCTGCCGCCCCCAGCCCCGACCCCTCGGCGCCCTCCAGACCTGGCGGCCCTGCCAAGC

TCACGGGAAGCTCCTGCCGCCCCCAGCCCCGACCCCTCGGCGCCCTCCAGACCTGGCGGCCCTGCCAAGC

GCGATGGGGGGTGCGGGGGCGTGCGGGGGGGCGGCGCGACCTGGCGGCGGCGGTCACGGGCCCCGTGCCT

GCGATGGGGGGTGCGGGGGCGTGCGGGGGGGCGGCGCGACCTGGCGGCGGCGGTCACGGGCCCCGTGCCT

CCGTAGGTCTGCGAGGAACAGAAGTGCGAGGAGGAGGTCTTCCCGCTGGCCATGAACTACCTGGACCGCT

CCGTAGGTCTGCGAGGAACAGAAGTGCGAGGAGGAGGTCTTCCCGCTGGCCATGAACTACCTGGACCGCT

TCCTGTCGCTGGAGCCCGTGAAAAAGAGCCGCCTGCAGCTGCTGGGGGCCACTTGCATGTTCGTGGCCTC

TCCTGTCGCTGGAGCCCGTGAAAAAGAGCCGCCTGCAGCTGCTGGGGGCCACTTGCATGTTCGTGGCCTC

TAAGATGAAGGAGACCATCCCCCTGACGGCCGAGAAGCTGTGCATCTACACCGACAACTCCATCCGGCCC

TAAGATGAAGGAGACCATCCCCCTGACGGCCGAGAAGCTGTGCATCTACACCGACAACTCCATCCGGCCC

GAGGAGCTGCTGGTAACCACTGGACCCCGCCGCCCCCCGCCCCCCGCGAGCCGCACGCAGGACCACGGGG

GAGGAGCTGCTGGTAACCACTGGACCCCGCCGCCCCCCGCCCCCCGCGAGCCGCACGCAGGACCACGGGG

CCGGGGAAGGTGCAGGCGGTGGCGGCCGGCCCGCCTCTGACATATCTGCTCCTCCGAGGGAGGGCGGCCC

CCGGGGAAGGTGCAGGCGGTGGCGGCCGGCCCGCCTCTGACATATCTGCTCCTCCGAGGGAGGGCGGCCC

CGCCGCCGGGCGTCCCTGTCCGGGGAGCGGGCGGGATCCTAGCCGCCCTCGTCCCGCCGCCCTGTGTGCG

CGCCGCCGGGCGTCCCTGTCCGGGGAGCGGGCGGGATCCTAGCCGCCCTCGTCCCGCCGCCCTGTGTGCG

CTTGCCTGCGACTCCCACCGCGTTCGCGCCCCGCGGTGTGGCCGAAAAGTGGGCGGCGCGCGCCCTCCAG

CTTGCCTGCGACTCCCACCGCGTTCGCGCCCCGCGGTGTGGCCGAAAAGTGGGCGGCGCGCGCCCTCCAG

CGGCTGCACGAGGAGCGCCGCGCTCGGCGCTGAGCCTCCAGTTCCAGGTGGTGGGAGGTCTTTTTGTTTC

CGGCTGCACGAGGAGCGCCGCGCTCGGCGCTGAGCCTCCAGTTCCAGGTGGTGGGAGGTCTTTTTGTTTC

CACTTGCAGAGTCTTTTCACGCGGCGGGCGCCTTTTCTGTTTTGATCTGGGATTGCGTGTTGCCCCAGCT

CACTTGCAGAGTCTTTTCACGCGGCGGGCGCCTTTTCTGTTTTGATCTGGGATTGCGTGTTGCCCCAGCT

CCCTTGAGTCCCCAGCATTCGCCAGCCCTCCCCTCCAACATCCAGGACCGCACGAGACGCAGGGGCCAGT

CCCTTGAGTCCCCAGCATTCGCCAGCCCTCCCCTCCAACATCCAGGACCGCACGAGACGCAGGGGCCAGT

GCTCTGAGCCGGAGGTGCGGCGTGGCCCGGCCCCCGTGCTGCCGGCTTCCCCGCGCCCCCGGGCTGGCCC

GCTCTGAGCCGGAGGTGCGGCGTGGCCCGGCCCCCGTGCTGCCGGCTTCCCCGCGCCCCCGGGCTGGCCC

GCACCTCCCCTGATGGCCGCTCACCCTGTGTTCGCAGCAAATGGAGCTGCTCCTGGTGAACAAGCTCAAG

GCACCTCCCCTGATGGCCGCTCACCCTGTGTTCGCAGCAAATGGAGCTGCTCCTGGTGAACAAGCTCAAG

TGGAACCTGGCCGCAATGACCCCGCACGATTTCATTGAACACTTCCTCTCCAAAATGCCAGAGGCGGAGG

TGGAACCTGGCCGCAATGACCCCGCACGATTTCATTGAACACTTCCTCTCCAAAATGCCAGAGGCGGAGG

AGAACAAACAGATCATCCGCAAACACGCGCAGACCTTCGTTGCCCTCTGTGCCACAGGTAGGGCAGGCCC

AGAACAAACAGATCATCCGCAAACACGCGCAGACCTTCGTTGCCCTCTGTGCCACAGGTAGGGCAGGCCC

GGCAGCCCCCGGCCTCCCCTTGAGAGCCGGCTCCTTAGGTGACCCTGGCCGGCTTCTTGCTCTCCACCTG

GGCAGCCCCCGGCCTCCCCTTGAGAGCCGGCTCCTTAGGTGACCCTGGCCGGCTTCTTGCTCTCCACCTG

GGTGCTGTCTGGGAAGATGTCCCCAGACCCCCTCCTGCGCTGGAGAGCGCTCTTCCAGCTCTGGTGAGCA

GGTGCTGTCTGGGAAGATGTCCCCAGACCCCCTCCTGCGCTGGAGAGCGCTCTTCCAGCTCTGGTGAGCA

GAGGCCCTGGATTGTTTGTCGCGCTGGATGGAGGGAGATTTGCTCCCTCACGGCCACCATGCAGTACCTT

GAGGCCCTGGATTGTTTGTCGCGCTGGATGGAGGGAGATTTGCTCCCTCACGGCCACCATGCAGTACCTT

GGGCATTGGTGTGGACGGCTCAGCCTGCCTGTGTCCCGTTACTCTGGCCTCGTCCTTCAGGCCAGGCAGC

GGGCATTGGTGTGGACGGCTCAGCCTGCCTGTGTCCCGTTACTCTGGCCTCGTCCTTCAGGCCAGGCAGC

CTGTGGCCACTCCATGCTGAAAGGGGTTTACCTTGGCCACAGGGCCGCCTCCTTTCTCCACCCACCTCCA

CTGTGGCCACTCCATGCTGAAAGGGGTTTACCTTGGCCACAGGGCCGCCTCCTTTCTCCACCCACCTCCA

GCCCTTCTTGTGTCCTTAAGGAGCCTGAGCTGCAGAGGCCCCCTCCTGGCCTCTCCCAGGCTGGGCCACC

GCCCTTCTTGTGTCCTTAAGGAGCCTGAGCTGCAGAGGCCCCCTCCTGGCCTCTCCCAGGCTGGGCCACC

TGCCAGAGGCGCCTCCAGGGGCGGGGAGAGCTGTCGGCCTGCCTGCACCACGTGCTCTGGGCAGCCGAGT

TGCCAGAGGCGCCTCCAGGGGCGGGGAGAGCTGTCGGCCTGCCTGCACCACGTGCTCTGGGCAGCCGAGT

GCAGGGGTGTCCAGCAGAGGAGCTCGGCTGCCTGAGGCCCTGCCAGGGGTGCCGGCAGCCAGCCGGGCTC

GCAGGGGTGTCCAGCAGAGGAGCTCGGCTGCCTGAGGCCCTGCCAGGGGTGCCGGCAGCCAGCCGGGCTC

AGCTGAGCCCTGAGGGGGCGCTTCAGAGCACTCTCAGCTTGGGCCGCCACCGTGGGCAGCAGAAGCACCC

AGCTGAGCCCTGAGGGGGCGCTTCAGAGCACTCTCAGCTTGGGCCGCCACCGTGGGCAGCAGAAGCACCC

AGTCCTCACTTCCCCTGGCATGGCCCCAGAGGCCCCTCCCTGACATGGCCTTGGCCCCAGAACCCAGTGG

AGTCCTCACTTCCCCTGGCATGGCCCCAGAGGCCCCTCCCTGACATGGCCTTGGCCCCAGAACCCAGTGG

GGACAGACTCGCACATACACAGGGTGCCGCCTCCTGCTGTCCCCAGCCCTGCCTCTGACCCCCCTGTGAC

GGACAGACTCGCACATACACAGGGTGCCGCCTCCTGCTGTCCCCAGCCCTGCCTCTGACCCCCCTGTGAC

CGCCTCCTTCCCTGGCCCAGGAGGCCTGGTTACCTTCATGGGGGAGCATGGCCCCATCCCACCCAGCTCT

CGCCTCCTTCCCTGGCCCAGGAGGCCTGGTTACCTTCATGGGGGAGCATGGCCCCATCCCACCCAGCTCT

GCTGTGGCCCACCTTTGGTCAAGCCTCAGTTGTCACATCTGTTTGGGGGCTCACTCTGGGTGACCTAGGC

GCTGTGGCCCACCTTTGGTCAAGCCTCAGTTGTCACATCTGTTTGGGGGCTCACTCTGGGTGACCTAGGC

CACAAGGCCCACGGGGCATCAAAGAGGCAGTAGCATCTTCTCCCCTCCCCAGAGGGCAGAGCCCCCCAAG

CACAAGGCCCACGGGGCATCAAAGAGGCAGTAGCATCTTCTCCCCTCCCCAGAGGGCAGAGCCCCCCAAG

CCTACTTCAGAGCTCCCTTCTGACACCGGTAGCCCGCAGCCGGTATTCCAGAATGGGTTCTGGTTTAGGC

CCTACTTCAGAGCTCCCTTCTGACACCGGTAGCCCGCAGCCGGTATTCCAGAATGGGTTCTGGTTTAGGC

GTGAGGCCTCCCCCACCTCCTCCACCTGCTTGGGGCATGAACCCCTCCCCCACGTTTCCAAGCGAGTCCC

GTGAGGCCTCCCCCACCTCCTCCACCTGCTTGGGGCATGAACCCCTCCCCCACGTTTCCAAGCGAGTCCC

CAAGGTGGGCAGATGAAGATGCCAAGGATGTCGACCAGTCTGGATGGGTCTGGGGTGGGGGGGCATGCGG

CAAGGTGGGCAGATGAAGATGCCAAGGATGTCGACCAGTCTGGATGGGTCTGGGGTGGGGGGGCATGCGG

CAGACAGGGAGGCATTCTCTGGCTGGTGCTCCTCAGAGGAGAGAGGCCTCCGGAGACTCCAGACAGCCTT

CAGACAGGGAGGCATTCTCTGGCTGGTGCTCCTCAGAGGAGAGAGGCCTCCGGAGACTCCAGACAGCCTT

TTATGGAGCTGAAAGTGGCTTCAGAGAAATGCAAAGTTTCCTGGAGAGAACGTGGGGCGTGGTTCTTGCA

TTATGGAGCTGAAAGTGGCTTCAGAGAAATGCAAAGTTTCCTGGAGAGAACGTGGGGCGTGGTTCTTGCA

CAGCCTCCCTACAGGGTGGCTCCAGCAGTGGAGCTCCCCTCCCAGGACCCCTGGGTGCTAGTGGGAGGCA

CAGCCTCCCTACAGGGTGGCTCCAGCAGTGGAGCTCCCCTCCCAGGACCCCTGGGTGCTAGTGGGAGGCA

GTGGGCAGGTGCAGATTCTCGTCCTTCCCACTACTGCACACCCTTTGTCTGCGAAGGCGCCCCCAGCGGT

GTGGGCAGGTGCAGATTCTCGTCCTTCCCACTACTGCACACCCTTTGTCTGCGAAGGCGCCCCCAGCGGT

GGGTGAAGGAGGAGGGACACTTGGGGACCCAGCTGTGCACGTGCTCTCAGTGACTGTGGAGTCCACTCCA

GGGTGAAGGAGGAGGGACACTTGGGGACCCAGCTGTGCACGTGCTCTCAGTGACTGTGGAGTCCACTCCA

GGGTGGGTCCCGAGGGAGGGGCAGGAGACCAGGGGACCCACCCCTGCAAAGTGCTCCGGGTCCTGACCCG

GGGTGGGTCCCGAGGGAGGGGCAGGAGACCAGGGGACCCACCCCTGCAAAGTGCTCCGGGTCCTGACCCG

TGGCCACCCCATGGAACGTAACTGAGCAGCCAGTGCCTTGTTCCTGCTGGACATCTGTGGAGACAAGAGT

TGGCCACCCCATGGAACGTAACTGAGCAGCCAGTGCCTTGTTCCTGCTGGACATCTGTGGAGACAAGAG

GACTTACGGCTGCTTAAAGTCAGAAACAGGTTGAAGGAGGTGGAGGCGTGGGAAAGAGTCTAGGAAGGTG

GACTTACGGCTGCTTAAAGTCAGAAACAGGTTGAAGGAGGTGGAGGCGTGGGAAAGAGTCTAGGAAGGTG

TTTTTGCCCTCCACGTGGCAAAGGTTACATTTAAAGGTGATGCTGGGTGTTCTCCCTGCACTAGGCATTC

TTTTTGCCCTCCACGTGGCAAAGGTTACATTTAAAGGTGATGCTGGGTGTTCTCCCTGCACTAGGCATTC

CTGGCCCCAGGTCCCCAGCAGGTGTGCACATGCTGCATACACTCACGCATGGGGGTTTCAGGGCAGGTGC

CTGGCCCCAGGTCCCCAGCAGGTGTGCACATGCTGCATACACTCACGCATGGGGGTTTCAGGGCAGGTGC

GCCCTTGGCTCCGTGGGAGGCCAGGTGAGGAACGTCCAGTGCCAAGGAGCTTCCGGGACAGCTGTCACTT

GCCCTTGGCTCCGTGGGAGGCCAGGTGAGGAACGTCCAGTGCCAAGGAGCTTCCGGGACAGCTGTCACTT

CCCTTTACAACCAGGCAGCGGATAGGGTCAAATCCTGGAGCTTTGGTGTCTAATTCTGGGTGGCTCCTAA

CCCTTTACAACCAGGCAGCGGATAGGGTCAAATCCTGGAGCTTTGGTGTCTAATTCTGGGTGGCTCCTAA

TCTAAGCACAGACAGCACCACACACTGGGGTGGGGGCACGAGCTTCTGAAACAACGTGGCCCCAGTGACT

TCTAAGCACAGACAGCACCACACACTGGGGTGGGGGCACGAGCTTCTGAAACAACGTGGCCCCAGTGACT

CCACGCTGTGTGTGCCCCTGGAGACGGGGGGGTGCACAAGGTGCGGAGCCAGCTAGAACCTGTCGCTCCC

CCACGCTGTGTGTGCCCCTGGAGACGGGGGGGTGCACAAGGTGCGGAGCCAGCTAGAACCTGTCGCTCCC

TGCAGAAGCGGTTTCTGTGTGCGGTTCTGATTTGCCTCAATGAGAAGGTTTTCATTCATGGCTCCCGGCT

TGCAGAAGCGGTTTCTGTGTGCGGTTCTGATTTGCCTCAATGAGAAGGTTTTCATTCATGGCTCCCGGCT

CTCAGACTGGGTGGAACTGCTCCCATTTAAAGGGGAAAAGAGGTGGCTCGGCTCGTTAAGGATTTCTTTT

CTCAGACTGGGTGGAACTGCTCCCATTTAAAGGGGAAAAGAGGTGGCTCGGCTCGTTAAGGATTTCTTTT

TCTAAGTTGTTACGGCGCCCAGCAGCCGGCTTTGTCTCCCCTTCAGGGTGGCTGCCTTTCTTCCCGGCCC

TCTAAGTTGTTACGGCGCCCAGCAGCCGGCTTTGTCTCCCCTTCAGGGTGGCTGCCTTTCTTCCCGGCCC

CTCGCCGGCGGCCCTCTCTTTAACAAGGCCGAAGTTGTTTATTCTCTCGGGATGAAGTCTCGGATGGGCC

CTCGCCGGCGGCCCTCTCTTTAACAAGGCCGAAGTTGTTTATTCTCTCGGGATGAAGTCTCGGATGGGCC

GCCACACCCCTGGCGGCCCGTGGGGGCCCCTCTCCCTTTGTGCCTGGGTCGGCTCCCATTCAGCTCCCCC

GCCACACCCCTGGCGGCCCGTGGGGGCCCCTCTCCCTTTGTGCCTGGGTCGGCTCCCATTCAGCTCCCCC

GACCCCCCTTGTTCCCGGGCGCTCAGTGGCGCGAGATGAGGCGATGGGGCCGACAAAGATGCCACACTCA

GACCCCCCTTGTTCCCGGGCGCTCAGTGGCGCGAGATGAGGCGATGGGGCCGACAAAGATGCCACACTCA

TCCCTGCCGACGTCCGGCTCCCAGCCCAGGGCCCCTGGTTCCTGTGCAGAATTCCTCGTGGGTGTGACAA

TCCCTGCCGACGTCCGGCTCCCAGCCCAGGGCCCCTGGTTCCTGTGCAGAATTCCTCGTGGGTGTGACAA

AAGGCTGCCCCCAGGCTCCGCTGGGGTGGGGGCCAGGCCAAGAGGCACATCCCACACTGGCCCACCTGTC

AAGGCTGCCCCCAGGCTCCGCTGGGGTGGGGGCCAGGCCAAGAGGCACATCCCACACTGGCCCACCTGTC

CACGGTAGGCGCATGACTGCCCTGAGGAGGGGAGGCCGGCATTCCCCGCCACAAACCAGGACGTAATTGG

CACGGTAGGCGCATGACTGCCCTGAGGAGGGGAGGCCGGCATTCCCCGCCACAAACCAGGACGTAATTGG

TGGCAGGGCTCTCTGTGGAAAGAGCCAGTCTGCTGTTTGTCTAGGAGGTCAGTCACAGAGGCCCCGAGAC

TGGCAGGGCTCTCTGTGGAAAGAGCCAGTCTGCTGTTTGTCTAGGAGGTCAGTCACAGAGGCCCCGAGAC

GCCCACTACTGCAGCCTGGCAGGCGGATGAGCCCAGTATCTGGCAGTGACCAGAGGGAGTTTTGTGCAGA

GCCCACTACTGCAGCCTGGCAGGCGGATGAGCCCAGTATCTGGCAGTGACCAGAGGGAGTTTTGTGCAGA

CCACAAAGGCTGATGGGCCGCCCTAGATTGGTGTCCCTCTTGGAAGTGGGCCCAGATGTGCGGGACAGTC

CCACAAAGGCTGATGGGCCGCCCTAGATTGGTGTCCCTCTTGGAAGTGGGCCCAGATGTGCGGGACAGTC

CCCAGGAAGCCCCAGGTGAGGGCACTGGTGCCCTCTTGGGAAAGCTGCTCCCTCCTGGGGCCCGGCTCCC

CCCAGGAAGCCCCAGGTGAGGGCACTGGTGCCCTCTTGGGAAAGCTGCTCCCTCCTGGGGCCCGGCTCCC

GGCCCAGTCCTCCAGGGGTGTCCCATGGTGACTGGTGCTAGGAACCCCACACCTCTTCCCTTACTTGGGA

GGCCCAGTCCTCCAGGGGTGTCCCATGGTGACTGGTGCTAGGAACCCCACACCTCTTCCCTTACTTGGGA

AGTCACTGGAATTGTTGGGCTACATCAGACGGCCCAGAAAAGTGTTTTTGTCATCGGCCAGAAATAGGAG

AGTCACTGGAATTGTTGGGCTACATCAGACGGCCCAGAAAAGTGTTTTTGTCATCGGCCAGAAATAGGAG

AGTTGTGAGTAGAGGGCCCGGGTGGAGTTGGGGTGTACTTGGTCTGTGCTCTGAAGGTCACTGTGACAGT

AGTTGTGAGTAGAGGGCCCGGGTGGAGTTGGGGTGTACTTGGTCTGTGCTCTGAAGGTCACTGTGACAGT

CATGGTCCCATGGTAAGGGGCATGGGTTGCTGGAAGAGCTCTTCCTTCCCGAGTGAGCCAAGCCGGGCTC

CATGGTCCCATGGTAAGGGGCATGGGTTGCTGGAAGAGCTCTTCCTTCCCGAGTGAGCCAAGCCGGGCTC

TCCTGGCGCCAGGGCCTGAGCCGCAGCCACACCACAGCCGCCCTGAAGGCTGCCGGCCAGGGCTTACCCC

TCCTGGCGCCAGGGCCTGAGCCGCAGCCACACCACAGCCGCCCTGAAGGCTGCCGGCCAGGGCTTACCCC

TCAAGGGACACGGAATGGCTTCATCAGTACCCTGCAGCCCCGTGGCCTGGCCCGGGTGGAGGCCTAGGCT

TCAAGGGACACGGAATGGCTTCATCAGTACCCTGCAGCCCCGTGGCCTGGCCCGGGTGGAGGCCTAGGCT

TCAGCCATGCGATGTCCCTTCAGAATATGACTTGTCTGCAATCCCTGCTGCTGGGGGGTGGCAGGTACTT

TCAGCCATGCGATGTCCCTTCAGAATATGACTTGTCTGCAATCCCTGCTGCTGGGGGGTGGCAGGTACTT

GGGGTGAGGGTTAGGGTCATAGAAGCGACATCTCTACGTCCTCATATTTGCGTCATCTAATTTTGTTTTT

GGGGTGAGGGTTAGGGTCATAGAAGCGACATCTCTACGTCCTCATATTTGCGTCATCTAATTTTGTTTTT

GTGAATACGTGATAACATTCACAAGGCTCAAGATGCTAAAAGGATGAGAAGGCAGTGATGTCCCCATCAC

GTGAATACGTGATAACATTCACAAGGCTCAAGATGCTAAAAGGATGAGAAGGCAGTGATGTCCCCATCAC

CTGTCCTGTGTCTTCCCGTGGCTTTCTCTTTCCTTGGTTATGTTTGAGTCAACAGTGGGGCTGACGTTCC

CTGTCCTGTGTCTTCCCGTGGCTTTCTCTTTCCTTGGTTATGTTTGAGTCAACAGTGGGGCTGACGTTCC

AGGAGGGTCCGTGGGCCAGGCTCTTGCTCTCCGAGTGCCCAGGGATGGCTGGAGGCTGAGGAGGGCCTGG

AGGAGGGTCCGTGGGCCAGGCTCTTGCTCTCCGAGTGCCCAGGGATGGCTGGAGGCTGAGGAGGGCCTGG

ATGTGGAGCCTCAGATACCGAGTGCTTCCCTTCAGGCCGGGCCGCTTGCTCAGAGCCAGCACACAGGGAT

ATGTGGAGCCTCAGATACCGAGTGCTTCCCTTCAGGCCGGGCCGCTTGCTCAGAGCCAGCACACAGGGAT

GCCCGGATCACGGGGGCCCTGAGAGGGTCCCCTGCTCACAGCCTCCTTCCCTCTCTCCTTCTGCCTCAGA

GCCCGGATCACGGGGGCCCTGAGAGGGTCCCCTGCTCACAGCCTCCTTCCCTCTCTCCTTCTGCCTCAGA

TGTGAAGTTCATTTCCAATCCGCCCTCCATGGTGGCAGCGGGGAGCGTGGTGGCCGCAGTGCAAGGCCTG

TGTGAAGTTCATTTCCAATCCGCCCTCCATGGTGGCAGCGGGGAGCGTGGTGGCCGCAGTGCAAGGCCTG

AACCTGAGGAGCCCCAACAACTTCCTGTCCTACTACCGCCTCACACGCTTCCTCTCCAGAGTGATCAAGT

AACCTGAGGAGCCCCAACAACTTCCTGTCCTACTACCGCCTCACACGCTTCCTCTCCAGAGTGATCAAGT

GTGACCCGGTAAGTGAGGGTGATGTCCCAGGCAGCCTTGCCGGGGCTTACAGGGGGAGACACCTAGTGCC

GTGACCCGGTAAGTGAGGGTGATGTCCCAGGCAGCCTTGCCGGGGCTTACAGGGGGAGACACCTAGTGCC

ACGGAAATGCCGAGGCTGGTGCCAAGGCCCCCAAGGGTGACAAGGTTGGGGCTGGGGCTGGGCCCCTCGG

ACGGAAATGCCGAGGCTGGTGCCAAGGCCCCCAAGGGTGACAAGGTTGGGGCTGGGGCTGGGCCCCTCGG

ACCCCAGGCCACAGACTGACAGGGCACCGGCTTCTTCCACTGCTCCTAGAACTTACTGACTGGCTGGGAG

ACCCCAGGCCACAGACTGACAGGGCACCGGCTTCTTCCACTGCTCCTAGAACTTACTGACTGGCTGGGAG

GTCCTCACAGCCTTCTCACGTCCCCTGGGGCTTCCAGGAGCCGTAGAGTTTCTGGGCGAAGCGTCCGGGA

GTCCTCACAGCCTTCTCACGTCCCCTGGGGCTTCCAGGAGCCGTAGAGTTTCTGGGCGAAGCGTCCGGGA

CGGAGGCCCCAGGCGGCCCCAGCCAATGGTCTGTGTGGTGATGGTGTGTGGGGTTAGGCCCAGGCGAGCT

CGGAGGCCCCAGGCGGCCCCAGCCAATGGTCTGTGTGGTGATGGTGTGTGGGGTTAGGCCCAGGCGAGCT

TTGTTTGGGCCACAATGTGCGTGGCCAATAAATAGATGCTTGAAAAGGGCTCCTGTGAGGTCCGAGACAC

TTGTTTGGGCCACAATGTGCGTGGCCAATAAATAGATGCTTGAAAAGGGCTCCTGTGAGGTCCGAGACAC

CGGACAACGGGCGGATAGAGACAGCCTTGTTGTTTACGGCCTCTTTGAGAGGCTGCTGCTGTTAAACCCT

CGGACAACGGGCGGATAGAGACAGCCTTGTTGTTTACGGCCTCTTTGAGAGGCTGCTGCTGTTAAACCCT

GGGATGACTGTGTCTTTCTTCTTAAAAATGCCATTGTTTTATTCCCGAGTCTTTTCTTAAAGAAAGAATT

GGGATGACTGTGTCTTTCTTCTTAAAAATGCCATTGTTTTATTCCCGAGTCTTTTCTTAAAGAAAGAATT

AAAATGACAATCAAAAGGGTTTGTGGCATTTACCAAATTAGACCAGAGAGGTGGCCGGGTCAGCCGCCGG

AAAATGACAATCAAAAGGGTTTGTGGCATTTACCAAATTAGACCAGAGAGGTGGCCGGGTCAGCCGCCGG

CCCCGCGGTGTGTGAGGGAGTGACCGCCTGACCCCAGCTTGGGGCTGGGTGGGCCTGCAAGACCCGTTTT

CCCCGCGGTGTGTGAGGGAGTGACCGCCTGACCCCAGCTTGGGGCTGGGTGGGCCTGCAAGACCCGTTTT

GGCTCTGGCCTGGGCCGCCTCTTGGTGGTCTGCCCTCGAGCCTCCCGGGGACTCCGCACGGGTCTCAGCA

GGCTCTGGCCTGGGCCGCCTCTTGGTGGTCTGCCCTCGAGCCTCCCGGGGACTCCGCACGGGTCTCAGCA

GATGCTATCTAGGGTCCACCTGCCTGTCCCCTGCCTAGTGGTGCCTCTGTCCCGGGGACACTGGGAGTAG

GATGCTATCTAGGGTCCACCTGCCTGTCCCCTGCCTAGTGGTGCCTCTGTCCCGGGGACACTGGGAGTAG

CGGCTGCCCAGCCCATGTGTGTCTCGGAAGAGGAAGAAGCTTTTTTGCCGTGGGACACCGAAGTTGGCAG

CGGCTGCCCAGCCCATGTGTGTCTCGGAAGAGGAAGAAGCTTTTTTGCCGTGGGACACCGAAGTTGGCAG

GGGCCTCCCTTCTGTGTTCTCGGCCATGGCCTCCCTTGCACCCTGCCCCGTGTTATCCTTTGGGGGTGGT

GGGCCTCCCTTCTGTGTTCTCGGCCATGGCCTCCCTTGCACCCTGCCCCGTGTTATCCTTTGGGGGTGGT

GAGGTGTCCTCACCCGCTGTAGGGTGGAGGCCAGCAGCCCGCAGCTCTCTCAGGAAAATGGCTCAGAAAC

GAGGTGTCCTCACCCGCTGTAGGGTGGAGGCCAGCAGCCCGCAGCTCTCTCAGGAAAATGGCTCAGAAAC

ACCATCGAGGCCTCCAGAAGCCCAGCAAAGAGAAAGCCCCTCCATCAAAATGAAACTCGCGTCTGCACTT

ACCATCGAGGCCTCCAGAAGCCCAGCAAAGAGAAAGCCCCTCCATCAAAATGAAACTCGCGTCTGCACTT

TTCATTTCGAACTCCACGCCCTGAGTGAAAACCGCTTCCCCGCCAGGGGTGACTGCCCTGGGATGTTGCT

TTCATTTCGAACTCCACGCCCTGAGTGAAAACCGCTTCCCCGCCAGGGGTGACTGCCCTGGGATGTTGCT

GTCTTCGGGCAGTTGTGGGAAGTTGGGCGCTGGCCCTTATTTGAGTAGAGACCATCTTAACTAGATTGGA

GTCTTCGGGCAGTTGTGGGAAGTTGGGCGCTGGCCCTTATTTGAGTAGAGACCATCTTAACTAGATTGGA

GGCACACGTCTCACAGCTGACAGACACACGGGGTGAAGTTACCCGAGGCGGAGTCCACTCTGCCTGATCA

GGCACACGTCTCACAGCTGACAGACACACGGGGTGAAGTTACCCGAGGCGGAGTCCACTCTGCCTGATCA

GCTAGTGACCAACGTAGCTGAGCCCAGACTCAGAAAAACCGTCCACAGCAGAGGCCCCTGCATTTTCTAG

GCTAGTGACCAACGTAGCTGAGCCCAGACTCAGAAAAACCGTCCACAGCAGAGGCCCCTGCATTTTCTAG

GGCGTGTTCTAGAATTTTCTTTGGTGGGTGGAATGTCCATCTGTGCAAATCGGGTGCGCAGTGCCACACA

GGCGTGTTCTAGAATTTTCTTTGGTGGGTGGAATGTCCATCTGTGCAAATCGGGTGCGCAGTGCCACACA

CCAGTGACTTTTCGCGGAGGAGCGTGCTGCCTTTTTGGAGCTTCTGGCTGTGGGAGAACAGCTTTGTCCA

CCAGTGACTTTTCGCGGAGGAGCGTGCTGCCTTTTTGGAGCTTCTGGCTGTGGGAGAACAGCTTTGTCCA

CCGGGGTAGCCTTGCAGGCAGCTGTGGGGCCAGAGGAATGAAGGAAGGTCCTGGAGTCTAGCTGCATGTG

CCGGGGTAGCCTTGCAGGCAGCTGTGGGGCCAGAGGAATGAAGGAAGGTCCTGGAGTCTAGCTGCATGTG

TGACCCTGGAGTGGGTCATGGGCGAGGGACGGGCCGCAGGTGAAGAATCCCTGGATGGAGCTGCCAGGCC

TGACCCTGGAGTGGGTCATGGGCGAGGGACGGGCCGCAGGTGAAGAATCCCTGGATGGAGCTGCCAGGCC

CCTGGGGCTGAGAATTGAAGCTGGCTGGTGTTTTAGGTTGAACGTCAGGAGTCTTGTATCTCACCCCAGG

CCTGGGGCTGAGAATTGAAGCTGGCTGGTGTTTTAGGTTGAACGTCAGGAGTCTTGTATCTCACCCCAGG

CCTCTGGCCTCAGTTTCCCCATCTGTACAGTGGGACTGTTTGTGCAGCCAGCCCGGCCAGCTTCATTTGC

CCTCTGGCCTCAGTTTCCCCATCTGTACAGTGGGACTGTTTGTGCAGCCAGCCCGGCCAGCTTCATTTGC

CATGATGAGAATTTATCTGAGGGGCGGGAGAGGAAAGCCCTCCCTATAAAGGTACAGGCGCTAAAATGTC

CATGATGAGAATTTATCTGAGGGGCGGGAGAGGAAAGCCCTCCCTATAAAGGTACAGGCGCTAAAATGTC

G*(101)*TGACCTCAGTGGTCCACCTAAAAGTCGTTCTGGCCTGGGTCATCGCCTGTCGTGCTATGCCTTT
GTCCA

A*(101)*TGACCTCAGTGGTCCACCTAAAAGTCGTTCTGGCCTGGGTCATCGCCTGTCGTGCTATGCCTTT
GTCCA

GCCCCTTCTGGTTGGGAGTTAAGTGGCACCTGTGCGGCACGTGGTGGGGCTGTGGCCCAGCCCTGCTCCT

GCCCCTTCTGGTTGGGAGTTAAGTGGCACCTGTGCGGCACGTGGTGGGGCTGTGGCCCAGCCCTGCTCCT

TGTGGAAGGTCTGTTTCCTGGGCTGCCTAGAGACTTGGCTTGAAGCCCTAGCGTGGCTTCCTGGCAGTTG

TGTGGAAGGTCTGTTTCCTGGGCTGCCTAGAGACTTGGCTTGAAGCCCTAGCGTGGCTTCCTGGCAGTTG

GGACACACACAGCCCCAACACATGGAGCCGGTTCTCCATCCAGAAGCCCCCGGGCAGTAAGCAGCCACTT

GGACACACACAGCCCCAACACATGGAGCCGGTTCTCCATCCAGAAGCCCCCGGGCAGTAAGCAGCCACTT

CAGGCTGCGTGGGACTTGCCCGTGGTGGAGCCTAGGAGAGGCCCCTGGCTGGGCGTGGCGTTCCAGATTT

CAGGCTGCGTGGGACTTGCCCGTGGTGGAGCCTAGGAGAGGCCCCTGGCTGGGCGTGGCGTTCCAGATTT

CACGGCTGCTCTTTCCCACTGACAGTGTGGTGTGGACGCTGCCAAGGGAGTCTGGAGCCCCAGAGGGTGG

CACGGCTGCTCTTTCCCACTGACAGTGTGGTGTGGACGCTGCCAAGGGAGTCTGGAGCCCCAGAGGGTGG

AGGTGCAGGACTTCCAGGAGCGTCCGTCGCACTCCACCCGAGGGCGAGCACCTCAGTGGCCGCAGTGGGT

AGGTGCAGGACTTCCAGGAGCGTCCGTCGCACTCCACCCGAGGGCGAGCACCTCAGTGGCCGCAGTGGGT

GGATGCATGCTGTGCCAGGCTGATGGCTGGCCCCGGGGCACAGGCCTGAGCGGGAGAGGATGGAGGGGAG

GGATGCATGCTGTGCCAGGCTGATGGCTGGCCCCGGGGCACAGGCCTGAGCGGGAGAGGATGGAGGGGAG

GGATCAATGGTCCAGGTCCCCCTGGCCACCCAGCATTCATCCTCAGTCATGCACGGCCCAAGGCTTCGAC

GGATCAATGGTCCAGGTCCCCCTGGCCACCCAGCATTCATCCTCAGTCATGCACGGCCCAAGGCTTCGAC

AGCCATTGATCATGGAAGGCCAGGTTCACCTCAAGGGCTGCCACATGGAGAGGTTAAGTCTGAAAAGGCT

AGCCATTGATCATGGAAGGCCAGGTTCACCTCAAGGGCTGCCACATGGAGAGGTTAAGTCTGAAAAGGCT

GAAAAGGCAGGGTTC *(102)* AAAGGGCCTCCTGTCCAGATCAGATGGCACTGAATTCCCCAGGGAGCTGG CACGG

GAAAAGGCAGGGTTA *(102)* AAAGGGCCTCCTGTCCAGATCAGATGGCACTGAATTCCCCAGGGAGCTGG CACGG

CCAGTGGGAACAGGCGGTGAAGGCGCTGTTGGACATGGGGACGGGCAGGGGGTGTGCAGGGTGGGCGGGC

CCAGTGGGAACAGGCGGTGAAGGCGCTGTTGGACATGGGGACGGGCAGGGGGTGTGCAGGGTGGGCGGGC

AAGCATCTGGTGTCTTGTGGCTCCAGAGACCAGGTGGGAGGTGGAGGCA *(103)* TTTGGTCCTGAGTGTC CTGAC

AAGCATCTGGTGTCTTGTGGCTCCAGAGACCAGGTGGGAGGTGGAGGCG *(103)* TTTGGTCCTGAGTGTC CTGAC

AGGTGATGGCAGCTCCCACATCTCGCTCAGGTTCAGAGGAGGCAGCATGGGCCGAGGGACAGTTTTTGGC

AGGTGATGGCAGCTCCCACATCTCGCTCAGGTTCAGAGGAGGCAGCATGGGCCGAGGGACAGTTTTTGGC

TTAGTCTTGCTCTTATAAAGGCTTCCGGGTCATGGCACCTGGGAAGGGGCCCTCGCTGCAGGCCCCTTCT

TTAGTCTTGCTCTTATAAAGGCTTCCGGGTCATGGCACCTGGGAAGGGGCCCTCGCTGCAGGCCCCTTCT

AAGGACCCCCTCTTCCCACCTCTCCCCACCCTCTCTCTCTCAGGACTGCCTCCGGGCCTGCCAGGAGCAG

AAGGACCCCCTCTTCCCACCTCTCCCCACCCTCTCTCTCTCAGGACTGCCTCCGGGCCTGCCAGGAGCAG

ATCGAAGCCCTGCTGGAGTCAAGCCTGCGCCAGGCCCAGCAGAACATGGACCCCAAGGCCGCCGAGGAGG

ATCGAAGCCCTGCTGGAGTCAAGCCTGCGCCAGGCCCAGCAGAACATGGACCCCAAGGCCGCCGAGGAGG

AGGAAGAGGAGGAGGAGGAGGTGGACCTGGCTTGCACACCCACCGACGTGCGGGACGTGGACATCTGAGG

AGGAAGAGGAGGAGGAGGAGGTGGACCTGGCTTGCACACCCACCGACGTGCGGGACGTGGACATCTGAGG

GCGCCAGGCAGGCGGGCGCCACCGCCACCCGCAGCGAGGGCGGAGCCGGCCCCAGGTGCTCCC *(104)* CT GACAG

GCGCCAGGCAGGCGGGCGCCACCGCCACCCGCAGCGAGGGCGGAGCCGGCCCCAGGTGCTCCA *(104)* CT GACAG

TCCCTCCTCTCCGGAGCATTTTGATACCAGAAGGGAAAGCTTCATTCTCCTTGTTGTTGGTTGTTTTTTC

TCCCTCCTCTCCGGAGCATTTTGATACCAGAAGGGAAAGCTTCATTCTCCTTGTTGTTGGTTGTTTTTTC

CTTTGCTCTTTCCCCCTTCCATCTCTGACTTAAGCAAAAGAAAAAGATTACCCAAAAACTGTCTTTAAAA

CTTTGCTCTTTCCCCCTTCCATCTCTGACTTAAGCAAAAGAAAAAGATTACCCAAAAACTGTCTTTAAAA

GAG(105)A GAGAGAGAAAAAAAAAAATAGTATTTGCATAACCCTGAGCGGTGGGGGAGGAGGGTTGTGC
TACAG

AAA(105)AAGAGAGAGAAAAAAAAAAATAGTATTTGC

ATGATAGAGGATTTTATACCCCAATAATCAACTCGTTTTTATATTAATGTACTTGTTTCTCTGTTGTAAG
AATAGGCATTAACACAAAGGAGGCGTCTCGGGAGAGGATTAGGTTCCATCCTTTACGTGTTTAAAAAAAA
GCATAAAAACATTTTAAAAACATAGAAAAATTCAGCAAACCATTTTTAAAGTAGAAGAGGGTTTTAGGTA
GAAAAACATATTCTTGTGCTTTTCCTGATAAAGCACAGCTGTAGTGGGGTTCTAGGCATCTCTGTACTTT
GCTTGCTCATATGCATGTAGTCACTTTATAAGTCATTGTATGTTATTATATTCCGTAGGTAGATGTGTAA
CCTCTTCACCTTATTCATGGCTGAAGTCACCTCTTGGTTACAGTAGCGTAGCGTGCCCGTGTGCATGTCC
TTTGCGCCTGTGACCACCACCCCAACAAACCATCCAGTGACAAACCATCCAGTGGAGGTTTGTCGGGCAC
CAGCCAGCGTAGCAGGGTCGGGAAAGGCCACCTGTCCCACTCCTACGATACGCTACTATAAAGAGAAGAC
GAAATAGTGACATAATATATTCTATTTTTATACTCTTCCTATTTTTGTAGTGACCTGTTTATGAGATGCT
GGTTTTCTACCCAACGGCCCTGCAGCCAGCTCACGTCCAGGTTCAACCCACAGCTACTTGGTTTGTGTTC
TTCTTCATATTCTAAAACCATTCCATTTCCAAGCACTTTCAG

**Data file S1. Genome sequencing of the Cyclin D1 locus in A375 cells after mCitrine-Cyclin D1 knock-in.** Aligned genome sequencing of CCND1 locus in wildtype A375 cells (reference sequence) highlighted in pale blue compared to A375 F4 mCitrine-CCND1 cells which remain unhighlighted. The start of mCitrine is denoted in yellow, the linker is denoted in green, the start of the CCND1 coding region is denoted in pink and the end of the CCND1 coding region is denoted in orange. Known SNPs in the CCND1 are denoted in red followed by the reference SNP (rs) report cited. Exons are underlined while introns are not underlined in the reference sequence.

### Supplemental Movie Figures:

**Movie S1**. **Correlating CDK2 activity sensor and mCitrine-Cyclin D1 levels in dabrafenib-treated A375 cell.** Representative live-cell movie of the mCitrine-Cyclin D1 A375 line treated with 1 μM dabrafenib. Movie starts 40 h after drug addition. Left: CDK2 activity sensor imaged in mCherry. Right: endogenous mCitrine-Cyclin D1 tag imaged in YFP. Images were acquired once every 15 minutes. Scale bar = 50 μm. The yellow arrow follows a single cell that goes on to escape from drug addition.
